# DNAAF9 (Shulin) and ARL3 regulate the transport and activation of ciliary Outer Dynein Arms (ODAs)

**DOI:** 10.1101/2024.08.15.608058

**Authors:** Karim Housseini B Issa, Muyang Ren, Bradley Burnet, Charlotte Melia, Kate Heesom, Girish R. Mali

## Abstract

Multiciliogenesis requires large-scale biosynthesis of motility-powering axonemal inner and outer dynein arm motors (IDA and ODA) prior to their intraflagellar transport (IFT) into cilia. ODAs are inhibited by the packaging chaperone Shulin during ciliogenesis in *T. thermophila.* How Shulin is released for ODAs to become active inside cilia remains unclear. We establish interactions between DNAAF9 (human Shulin) and mammalian ODA subunits, IFT proteins and the ciliary small GTPase ARL3 using proteomics and *in vitro* reconstitutions. Mutagenesis combined with biochemical and structural studies reveal that DNAAF9 and Shulin preferentially bind the active Arl3-GTP state. GTP-loaded Arl3 can access, bind and displace Shulin from the packaged ODA-Shulin complex. We propose that once the inhibited ODA complex enters growing cilia, Arl3-GTP displaces Shulin (DNAAF9) and sequesters it away from ODAs promoting activation of their motility specifically inside cilia.

## Introduction

Motile cilia are microtubule-based extensions found on eukaryotic cell surfaces. Coordinated ciliary motion aids the locomotion of unicellular organisms, propels gametes, and causes the essential flow of biological fluids over tissues in multicellular organisms. Ciliary beating is powered by axonemal outer dynein arm (ODA) motors and the waveform is modulated by inner dynein arms (IDAs)^1^. Ciliary dysmotility due to defects in ODAs causes primary ciliary dyskinesia (PCD), a severe human ciliopathy characterised by chronic respiratory symptoms^2^.

ODAs and IDAs are macromolecular complexes that undergo cytoplasmic pre-assembly^3^. Large quantities of pre-assembled dyneins need to be transported to growing cilia and due to their size they are thought to mainly rely on the intraflagellar transport (IFT) system to enter cilia^4–6^. Following ciliary entry and transport to the growing tips, ODAs bind specific docking sites on doublet microtubules^7^ (**Fig. 1a**). Throughout their trafficking from the cytoplasm to their final docking sites in cilia, ODAs need to be kept inactive to prevent off-target interactions. Tight control of ciliary targeting and the regulated activation of dyneins is particularly important in mammalian multiciliated cells such as human airway epithelial cells or ciliated protozoa such as *Tetrahymena thermophila* which deploy several 1000s of pre-assembled ODA and IDA motors into several growing motile cilia. Premature cytoplasmic activation in such systems would have particularly deleterious consequences for the cells survival.

**FIGURE 1.**
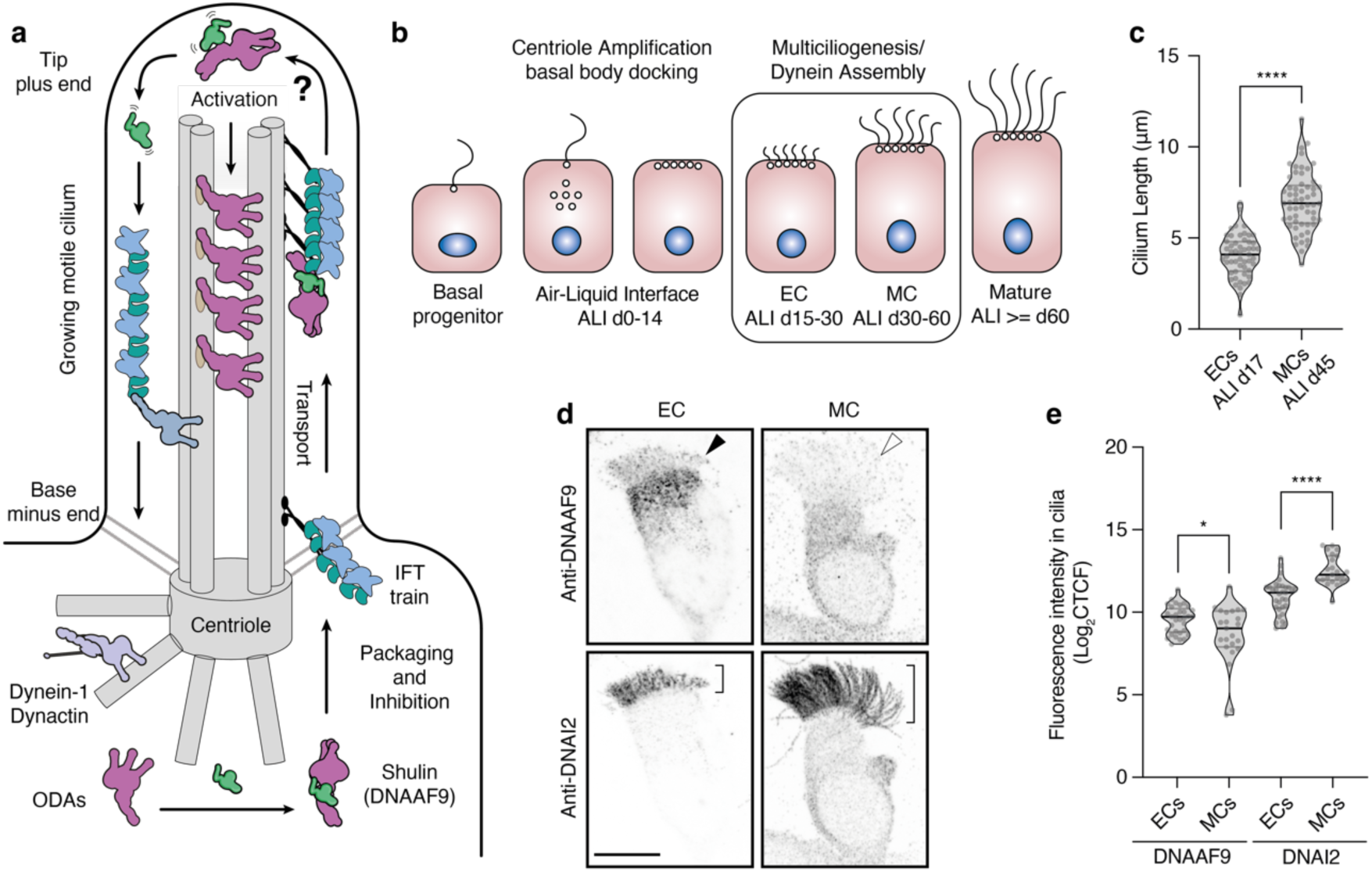
DNAAF9 enters growing motile cilia in differentiating human airway cells. **a.** Model of a motile cilium under construction involving Shulin/DNAAF9 for ODA packaging and inhibition. IFT trains transport ODAs to ciliary tips. How Shulin gets released for ODA activation inside cilia is an open question. **b.** Schematic of *in vitro* differentiation of human airway epithelial cells (HAECs) undergoing multiciliogenesis at the air-liquid interface (ALI). **c.** Cilia length measurements from ALI d17 (n=53 ECs) and ALI d45 (n=57 MCs) cells depicted as violin plots with the means shown as horizontal lines. **d.** Co-immunostaining of DNAAF9 and DNAI2 in an EC with short cilia (black arrowhead and short bracket) and an MC with long cilia (white arrowhead and long bracket). Scale bar, 10μm. **e.** Quantification of fluorescent signal intensities for DNAAF9 and DNAI2 in motile cilia of ECs (n=33) and MCs (n=23) depicted as violin plots with the means shown as horizontal lines. Unpaired (two-tailed) t-tests were used to calculate p-values; *p<0.05, ****p<0.0001 Data in (e) is log transformed (Log_2_[CTCF] = corrected total cilia fluorescence).

Motor inhibition of ODAs is achieved by the packaging chaperone Shulin which directly binds and locks them in a conformation that cannot productively engage with microtubules^8^. This inhibited ODA conformation is proposed to underpin their unimpeded ciliary import as it prevents aberrant interactions with cellular and ciliary microtubules during transport to ODA docking sites. The inhibited conformation of ODAs is similar to the compact inhibited conformation of dynein-2 on anterograde IFT trains^8, 9^. Shulin and ODAs co-localise in regenerating *Tetrahymena* cilia suggesting it remains bound to ODAs to dampen their motor activity during ciliary transport. There are several outstanding questions – is Shulin’s inhibitory mechanism conserved in its orthologs from other species such as its human ortholog DNAAF9?, does it interact with other transport factors to target packaged ODAs to cilia?, and most importantly, how is Shulin released from ODAs to allow their final activation inside cilia?

In this work we determine a key function of Shulin (DNAAF9) in targeting ODAs to cilia beyond its inhibitory role and present a molecular mechanism for ODA activation. DNAAF9 in human airway cells interacts with ODA complex subunits, IFT proteins and the ciliary small GTPase ARL3 which is essential for ciliogenesis and ciliary cargo trafficking. DNAAF9 binds and induces a closed conformation in ODAs purified from pig airway cilia. Furthermore, we show that Arl3 binds conserved surface exposed residues on the N1 domains of both DNAAF9 and Shulin in a GTP dependent manner. Modeling Arl3 binding at Shulin’s N1 domain and superimposing this model on the ODA-Shulin cryo-EM structure, highlights steric clashes, which we suggest, preclude the formation of a ternary complex between ODA, Shulin and Arl3. In support of this notion, Arl3-GTP displaces Shulin from an inhibited ODA-Shulin complex *in vitro*. Based on our data, we propose a model where the inhibition imposed by Shulin on ODAs is relieved by Arl3 in its active GTP state. As there are higher pools of active Arl3 reported inside cilia^10, 11^ the displacement of Shulin and the resulting activation of ODA motor activity would therefore occur in a spatially-restricted manner inside motile cilia.

## Results

### DNAAF9 enters growing airway multicilia and binds mammalian ODAs

Previously, Shulin’s human orthologue DNAAF9 (aka C20ORF194) was pulled-down with ARL3 which is a small GTPase that regulates the transport of both membrane and soluble ciliary cargo proteins^12–14^. This suggested to us that DNAAF9 could be involved in ciliary cargo transport of ODAs. We performed AlphaFold2 (AF2) analyses which revealed that DNAAF9 is structurally homologous to Shulin (**Fig. S1a, b, c, d, e**) – notably, sharing domains critical for binding and inhibiting ODAs (N1 domain and C3 extension) (**Fig. S1f, g, h**). Based on these observations, sequence conservation across evolution (**Fig. S2**) and previous findings, we reasoned that DNAAF9 could package ODAs and co-operate with ARL3 to regulate their ciliary trafficking in diverse species.

To verify functional conservation, we investigated DNAAF9’s functions using human airway epithelial cells (HAECs) cultured at the air-liquid interface (ALI). HAECs grown *in vitro* differentiate from lung basal progenitors into multiciliated cells after exposure to air (airlift). After initial stages of centriole amplification and basal body docking (ALI days 0-14), cells progress asynchronously from a basal state through early-mid (ALI days 15-30) and then mid-mature (ALI days 30-60) stages to reach full maturity (>60 days) (**Fig. 1b**). Ciliary lengths define each differentiation stage^15^. ALI cultures at earlier stages typically contain a mixed population of basal, early and mid-stage cells (here referred to as ECs) and start to grow short cilia (mean length 4.03 ± 1.17 μm from n=53 cells). Later ALI cultures contain a mixture of early, mid and mature-stage airway cells (here referred to as MCs) and on average have comparatively longer cilia (mean length 7.00 ± 1.6 μm from n=57 cells) (**Fig. 1c**). We reasoned that dynein assembly and trafficking would be most active at these stages of differentiation as cilia are still actively growing and have therefore investigated the role of DNAAF9 using ECs and MCs.

First, we tracked DNAAF9’s sub-cellular location in relation to the ODA intermediate chain DNAI2 which was used to mark cilia. Co-immunostaining ECs and MCs with anti-DNAAF9 and anti-DNAI2 antibodies (**Fig. 1c, d; Fig S3**) revealed DNAAF9 signal concentrated apically near basal bodies and was also observed in short growing multi-cilia of ECs (**Fig. 1d, e; Fig S3**). MCs showed more diffuse DNAAF9 signal in the cytoplasm and its levels were also slightly reduced from the longer MC cilia. This could indicate that most of the DNAAF9 gets retrieved from mature motile cilia for recycling or degradation in the cytoplasm (**Fig. 1d, e**). Importantly, DNAAF9’s localisation to growing human airway cilia resembles Shulin’s expression in *Tetrahymena* cells undergoing ciliary regeneration^8^. To gain further insights into DNAAF9’s role during airway cell differentiation we immunoprecipitated endogenous DNAAF9 from HAECs cultured to day 17 and day 45 post-airlift. Co-precipitating proteins were identified by tandem mass tag (TMT) labelling followed by mass spectrometry (MS) proteomics. DNAAF9 was the top hit from cells at both differentiation time points. We detected subunits of the human ODA holocomplex (DNAH5, DNAH9, DNAI1, LC3/NME9) amongst the proteins that co-precipitated with DNAAF9 (**Fig. 2a, b**). Western blot analysis of the IPs confirmed an interaction with the ODA intermediate chain DNAI2 (**Fig. 2c**).

**FIGURE 2.**
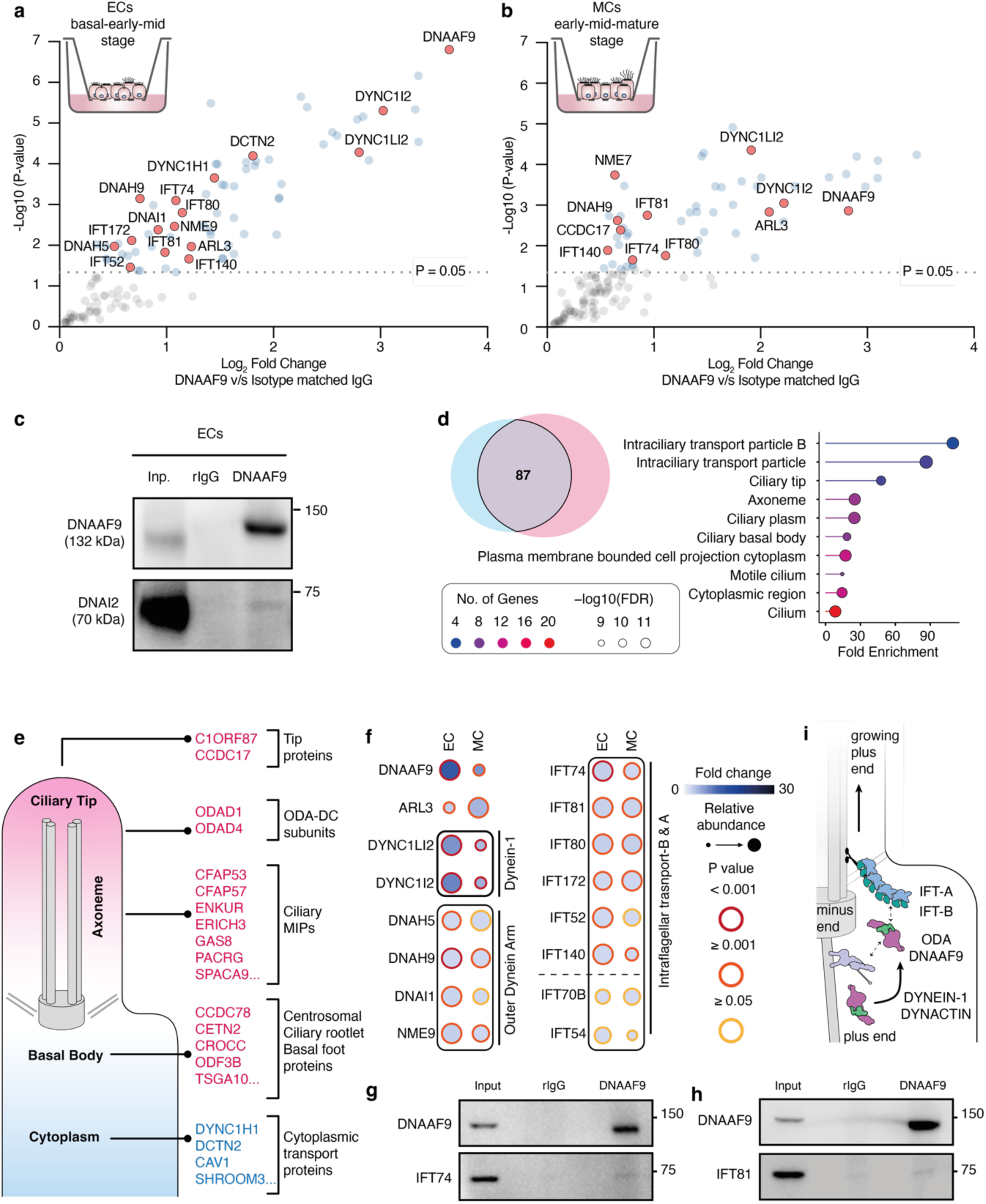
DNAAF9 interacts with ODA subunits and multiple transport proteins in human airway cells. **a., b.** DNAAF9 interactome from ALI d17 ECs and ALI d45 MCs. Proteins are plotted by enrichment (DNAAF9 IP vs isotype matched IgG IP) on the x-axis and significance (n=3 experimental replicates) on the y-axis. Proteins with p-values <0.05 (-log10 1.3) are colored in blue and key hits are labelled **c.** Co-IP blots for DNAAF9 and DNAI2 from ECs **d.** Venn diagram of 87 overlapping proteins from DNAAF9’s EC (blue) and MC (pink) interactomes. Dot plot shows gene ontology (GO) term enrichment of cellular component for the overlapping hits determined using ShinyGO with enriched terms (y-axis) ranked by fold enrichment (x-axis) over a background set of genes (FDR cutoff = 0.05). The points are coloured by number of proteins clustering within each term and sized by enrichment confidence (FDR) **e.** Proteins that exclusively co-precipitate with DNAAF9 either from ECs (in blue) or MCs (in pink) are grouped according to their sub-cellular location. **f.** Fold change and relative abundances of key DNAAF9 interactors from ECs and MCs are compared in a dot plot generated using ProHits-viz. The points are colored in shades of blue according to protein fold change and sized by relative abundance. The colour of the outline represents significance **g., h.** Co-IP blots for DNAAF9, IFT74 and IFT81 from ECs **i.** Schematic of a growing cilium summarising the key interactors of DNAAF9.

To verify the association of DNAAF9 with mammalian ODAs more directly, we purified ODAs from pig tracheal cilia using sucrose density gradient fractionation and verified the presence of ODA subunits co-sedimenting in the fractions by mass spectrometry (**Fig. S4**). Fraction 14 containing ODA heavy chains (HCs), intermediate chains (ICs) and light chains (LCs) (indicative of intact holocomplexes), was applied to an EM grid alone and after mixing with DNAAF9 and negatively stained. Intact double-headed ODA particles in isolation adopted the previously reported open ‘active’ conformation found in human respiratory cilia^16^ (**Fig. S5a**). Incubating DNAAF9 with fraction 14 causes a conformational change in pig ODAs from open to closed similar to the closed, inhibited ‘phi’-like state of triple headed *Tetrahymena* ODAs when bound by Shulin (**Fig. S5b, c**).

Shulin binds the *Tetrahymena* ODA γ-HC Dyh3 at two sites^8^. Shulin’s N1 domain contacts a conserved ‘LFGL’ motif in the γ-HC tail forming site 1. Shulin’s C-terminal alpha helical extension contacts conserved hydrophobic and acidic residues in γ-HCs motor near the AAA1(S) site. To model potential interactions between DNAAF9 and DNAH5 (human γ-HC) we performed AlphaFold3 (AF3) predictions using long sequences encompassing these sites. These structural predictions suggested potential contacts between DNAAF9s N2 domain and DNAH5’s ‘LFGL’ motif and between the C3 extension and conserved residues near the AAA1(S) site with moderate confidence (**Fig. S5d, e, f, g**). Taken together, these findings indicate that human DNAAF9 interacts with mammalian ODAs and maintains them in a closed conformation sharing its role as an inhibitor with its *Tetrahymena* ortholog Shulin.

### DNAAF9 interacts with proteins involved in cilia formation during airway cell maturation

In addition to ODA subunits, DNAAF9s EC and MC interactomes included the ciliary small GTPase ARL3, several IFT proteins and a few subunits of the cytoplasmic dynein-1 motor transport machinery (**Fig. 2a, b**). We performed Gene Ontology (GO) enrichment analysis to see if any proteins amongst the top 87 common hits in both the EC and MC interactomes clustered under specific GO terms. This highlighted an enrichment of IFT-B and ciliary tip proteins (**Fig. 2d**). We also identified proteins that uniquely co-precipitate with DNAAF9 in a cell stage-specific manner but at lower abundance. The DNAAF9 interactome from ECs (nb: mixture of basal, early and mid-cell stages) contained more cytoplasmic proteins (DYNC1H1, DCTN2, CAV1) in contrast to its interactome from MCs (nb: mixture of early, mid and mature-cell stages) which contained more centrosomal/basal body and axonemal proteins including ciliary tip proteins (C1ORF87, CCDC17), ODA docking complex subunits (ODAD1, ODAD4) and microtubule inner proteins (MIPs: CFAP57, ENKUR) (**Fig. 2e**)^17^. This observation suggests that DNAAF9’s interactions change during airway cell multiciliogenesis which could relate to its cellular functions in ciliary trafficking.

Using quantitative TMT proteomics, we next compared the relative abundances of top DNAAF9 protein interactors between ECs and MCs. This revealed that DNAAF9 levels reduce as airway cells mature which matches the overall reduction in DNAAF9 signals observed by immunostaining of MCs. This could be due to a downregulation in its gene expression or its enhanced protein degradation in the cell (**Fig. 2f**). DNAAF9 co-precipitates relatively higher levels of cytoplasmic dynein-1 light intermediate and intermediate chain subunits from ECs compared to MCs. Conversely, relatively higher levels of ARL3 co-precipitate with DNAAF9 from MCs compared to ECs (**Fig. 2f**). As several IFT-B subunits co-precipitated with DNAAF9 from both ECs and MCs (**Fig. 2b**) and IFT-B is a significantly enriched GO term, we verified two of the top IFT-B protein hits, IFT74 and IFT81, as novel interactors of DNAAF9 by IP western blot analyses (**Fig. 2g, h**). Based on the DNAAF9 protein interaction data, we propose the following sequence of events (**Fig. 2i):** First, DNAAF9 binds ODAs in the cytoplasm. Presently it is unclear if it binds ODAs co-translationally or after they are fully pre-assembled. ODAs are then transported to a peri-basal body pool likely via the dynein-1 transport machinery. This apically enriched pool of ODAs is maintained in an inhibited state by DNAAF9 to facilitate crossing the diffusion barrier at the transition zone via active transport on IFT trains.

### Arl3-GTP binds DNAAF9 *in vitro* and pulls-down Shulin from growing *Tetrahymena* cilia

Once inside cilia, DNAAF9/Shulin must detach from ODAs to allow their activation and stable axonemal incorporation for ciliary beating. Shulin’s release mechanism remains unknown. Based on coarse-grained MD-simulation of the ODA-Shulin complex, it has been proposed that conformational remodelling during ODA docking to doublet microtubules is sufficient to release Shulin^18^. However, these simulations were performed after decreasing the attractive forces between Shulin and ODA subunits by 0.3 times the default parameters to observe the detachment. We therefore reasoned that additional factors specifically inside cilia could first destabilize the ODA-Shulin complex triggering its release on microtubule association. As ARL3 is an established ciliary cargo release factor which interacts with DNAAF9 in airway cells we hypothesised that its binding to DNAAF9 could trigger its detachment from ODAs inside cilia. First, we probed this interaction by recombinantly expressing and reconstituting DNAAF9 with the *Tetrahymena* ARL3 orthologue (Arl3) in its GTP and GDP mimicking states using the variants Q70L and T30N respectively (**Fig. S6a**). In vertebrates, ARL3-Q71L is described as a dominant active variant due to defective GTP hydrolysis and conformationally it mimics the constitutive GTP-loaded active state. ARL3-T31N is the dominant negative variant which is defective in GTP binding and is thought to conformationally mimic the GDP-loaded inactive state^13, 19^. A pool of active ARL3 is maintained inside cilia due to the spatially restricted localisations of its GEF and GAP proteins, Arl13b on the ciliary membrane and Rp2 at the ciliary base respectively^20–22^. Reconstitutions indicate that only Arl3^Q70L^ co-elutes with DNAAF9 even in the presence of GDP (**Fig. 3a, b**) likely due to its stable GTP-locked conformation. All Arl3 variants co-elute with DNAAF9 in the presence of high GTP levels (**Fig. S6c, d**) which indicates that even the TN variant is able to bind DNAAF9 under saturating GTP conditions.

**FIGURE 3.**
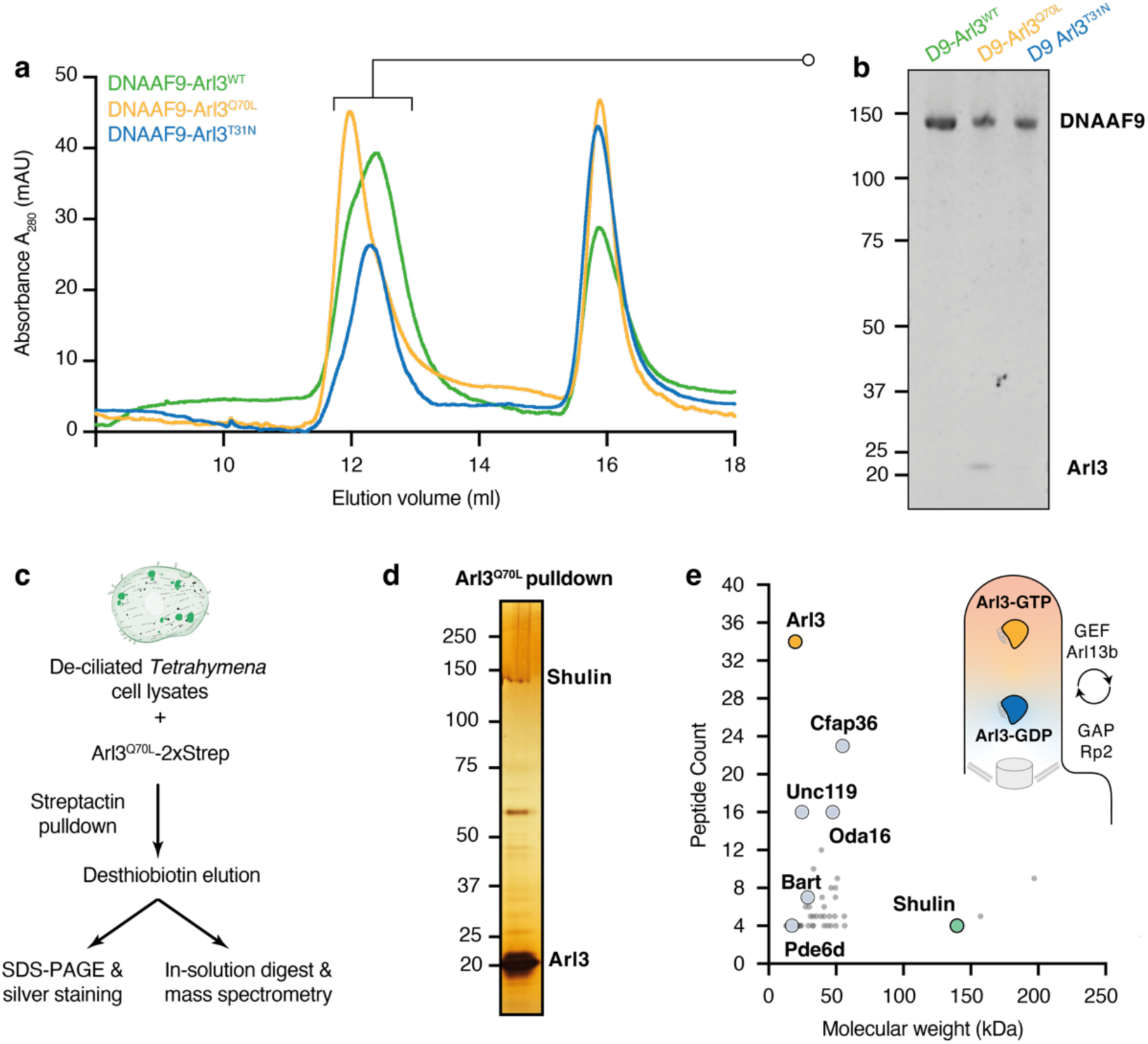
Arl3^Q70L^ co-elutes with DNAAF9 and pulls down endogenous Shulin from growing motile cilia. **a., b.** Analytical size-exclusion chromatography of DNAAF9-Arl3 complexes [three Arl3 variants: wildtype, GTP (Q70L) and GDP (T30N) mimicking mutants]. Proteins eluting in the first peak from each of the three reconstitutions are shown. **c.** Scheme used to identify interactors of activated Arl3 from deciliated *Tetrahymena* cells undergoing ciliary regeneration using the GTP-mimetic Q70L variant. **d.** Silver stained pull down gel showing proteins co-eluting with Arl3^Q70L^ **e.** Scatter plot shows proteins pulled down by Arl3^Q70L^ ranked by peptide count versus molecular weight. The schematic shows Arl3’s GTPase cycle. Active Arl3-GTP is enriched inside cilia by its ciliary GEF Arl13b and inactive Arl3-GDP is excluded as its GAP Rp2 localizes to the basal-body.

DNAAF9 was previously reported to co-purify with active ARL3-QL from primary ciliated human RPE1 cells^13^. Our proteomic data supports that both proteins also interact in differentiating human airway cells which have motile cilia. To test this interaction more directly in the context of actively ciliating cells, we performed a pull-down using recombinant *Tetrahymena* Arl3^Q70L^ as bait and lysates of *Tetrahymena* cells undergoing ciliary regeneration after deciliation (**Fig. 3c**). Analysis of the co-precipitating prey proteins identified several known effectors of Arl3 (Cfap36/Bartl1, Unc119, Pde6d)^21, 23^ and Shulin (**Fig. 3d, e**). We conclude that human DNAAF9 and *Tetrahymena* Shulin preferentially bind the GTP-mimicking active variant of *Tetrahymena* Arl3. Our findings, in the context of previous data, highlight the cross-species conservation of this interaction which occurs in both primary and motile ciliated cells. Overall, we establish DNAAF9 (Shulin) as a novel effector of ARL3 (Arl3) that functions under specific contexts such as during motile multiciliogenesis likely to regulate the transport of ODAs to cilia.

### ARL3 binds DNAAF9’s amino terminal domain

Next, we sought to visualize where ARL3 bound to DNAAF9 to ascertain whether its binding could explain DNAAF9’s, and also Shulin’s detachment from ODAs. We performed negative stain electron microscopy on purified recombinant DNAAF9 and Shulin in complex with Arl3^Q70L^. Single particle analysis provided well-defined 2D class averages (**Fig. 4a; S7b**). Comparing the averages with a calculated 2D projection profile of Shulin (PDB: 6ZYX_8) revealed the presence of an extra density adjacent to the amino-terminal N1 domain of DNAAF9/Shulin. This density was absent in about half of the class averages. We conclude that these class averages represent DNAAF9 and Shulin alone and in complex with Arl3 which binds the N1 domain.

**FIGURE 4.**
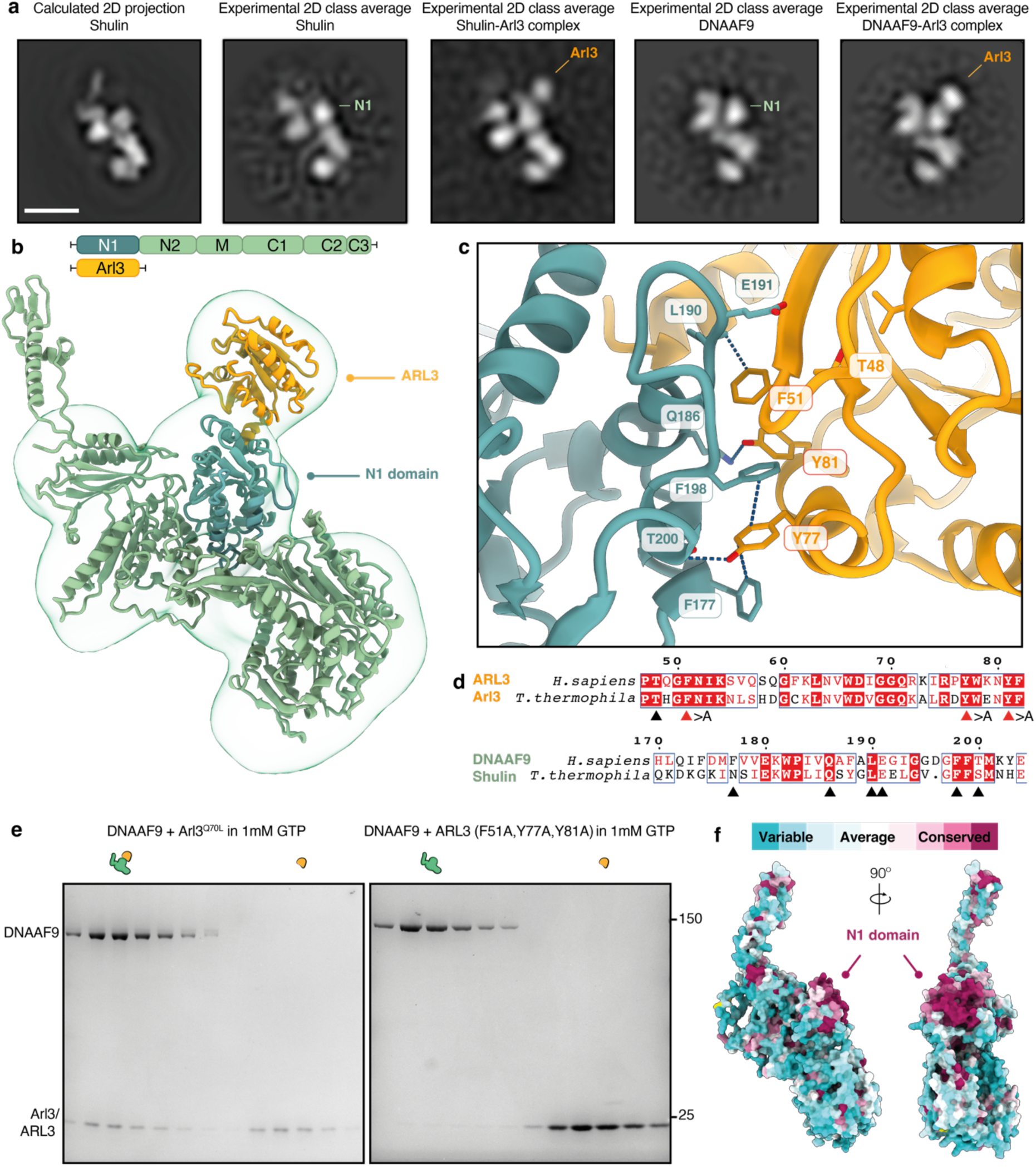
ARL3/Arl3 binds DNAAF9/Shulin at its conserved N1 domain via aromatic residues. **a.** Side view of a calculated 2D projection of Shulin (pdb:6ZYX_8) and best matching 2D class averages DNAAF9/Shulin alone and bound by Arl3^Q70L^. N1 domain and position of Arl3 are highlighted. Scale bar, 2.5 nm **b.** AF2-Multimer prediction of DNAAF9-ARL3 complex rigid-body docked into a negative stain EM density map of the complex. **c.** AF2-Multimer predicted binding interface between DNAAF9s N1 domain and ARL3 with residues forming non-covalent interactions (dashed bonds) labelled **d.** Sequence alignment of human and *Tetrahymena* DNAAF9/Shulin and ARL3/Arl3 with interface residues marked by arrowheads. Positions of single residues substituted with alanine (>A) residues for complex disruption are marked by red arrowheads. **e.** Analytical size-exclusion chromatography fractions from characterisations of complex formation between DNAAF9 and wildtype Arl3 or the triple alanine substitution ARL3 mutant. **f.** CONSURF scores mapped onto DNAAF9’s predicted structure. The N1 domain comprising the conserved surface patch where ARL3 binds is highlighted.

We next performed AF2-Multimer predictions to generate a structural model of the DNAAF9-ARL3 complex using full-length wildtype human sequences. The model predicted binding of ARL3 in a GTP-loaded conformation to the N1 domain of DNAAF9 in agreement with the 2D class averages and could be accommodated into an 18 Å 3D density map of the complex obtained by negative stain EM (**Fig. 4b**; **Fig. S7c, d, e**). Closer inspection of the binding region revealed a string of hydrogen bonds and π-π stacking interactions spanning the interface between conserved residues from DNAAF9s N1-domain and ARL3’s conserved amino terminal half (**Fig. 4c, d; Fig. S7f**). We reasoned that mutating interface residues in ARL3 could destabilise the complex and verify the predicted model. We generated an ARL3 triple mutant substituting three conserved interface aromatic residues (F51, Y77 and Y81) with alanine (**Fig. 4d)**. Reconstitutions under saturating GTP conditions showed that the interaction between DNAAF9/Shulin and the ARL3 triple mutant is disrupted compared to with the Arl3^Q70L^ GTP mimicking variant (**Fig. 4e; Fig. S8a**). We conclude that mutating the three aromatic residues destabilizes the interactions at the interface for the complex to be stably formed. Finally, we performed CONSURF analysis and mapped per-residue conservation scores on the predicted DNAAF9 structure. This revealed a highly conserved patch of surface exposed residues in the N1 domain in addition to a few conserved residues at the tip of the C3 extension that contact the motor domain (**Fig. 4f**). This analysis further supports the idea that the N1 domain harbours the binding site for ARL3 (Arl3) on DNAAF9 (Shulin).

To understand the nucleotide dependent binding of ARL3 to DNAAF9 we aligned our predicted model with previous X-ray structures of murine ARL3 in the GTP and GDP locked states (PDB: 4ZI2, 1FZQ respectively). Several structural features in ARL3 (α3 helix, interswitch beta strands β2-β3 and L3 loop) participate in the interaction with DNAAF9 (**Fig. S9a**). In the GDP-loaded state, the L3 loop and β2-strand reconfigure to form a β-hairpin which displaces the interacting α3 helix and sterically clashes with DNAAF9’s N1 domain (**Fig. S9b**). We also performed AF3-Multimer predictions between DNAAF9 and ARL3 to model the complex in the presence of GTP and GDP. AF3 generated a high confidence model for the interaction between GTP loaded ARL3 and DNAAF9 positioning it at the N1 interface but failed to model an interaction with GDP loaded ARL3 (as evidenced by a lack of residue level pair-wise association in the PAE plots) likely due to steric clashes (**Fig. S9c, d**). Overall, these structural analyses provide a molecular explanation of why only ARL3-GTP is able to bind DNAAF9.

### Active Arl3 binds and displaces Shulin from ODAs

Arl3 and ODAs contact Shulin’s N1 domain at distinct binding sites. Although there is no direct overlap between these sites, superimposing the Shulin-Arl3 prediction onto the cryo-EM guided structure of ODA-Shulin highlighted steric clashes between Arl3 and ODA heavy chains (**Fig. 5a, b, c**). As such clashes would either destabilise the ODA-Shulin complex or prevent stable complex formation, we hypothesised that Arl3 binding displaces Shulin from packaged (inhibited) ODAs and sequester it away from open (activated) ODAs to prevent rebinding. To test this, we first reconstituted ODAs purified from cilia with recombinant Shulin which induces the ‘phi’-like closed conformation^8^ (**Fig. 5d**; left-hand gel image). We then incubated the purified ODA-Shulin complex with an excess of Arl3^Q70L^ in the presence of GTP *in vitro* and resolved the resulting complexes by gel filtration. We observed low levels of Shulin co-eluting in fractions containing the ODA complex and most of the Shulin was found in complex with Arl3^Q70L^ (**Fig. 5d**; right-hand gel image). We conclude that adding Arl3 to the purified ODA-Shulin complex displaces most of the bound Shulin. Formation of a Shulin-Arl3 complex sequesters Shulin preventing it’s rebinding to ODAs. We propose that by displacing Shulin, Arl3 relieves the inhibition imposed on ODAs, allowing the motors to undergo structural reconfigurations required for the activation of motility (**Fig. 5e**).

**FIGURE 5.**
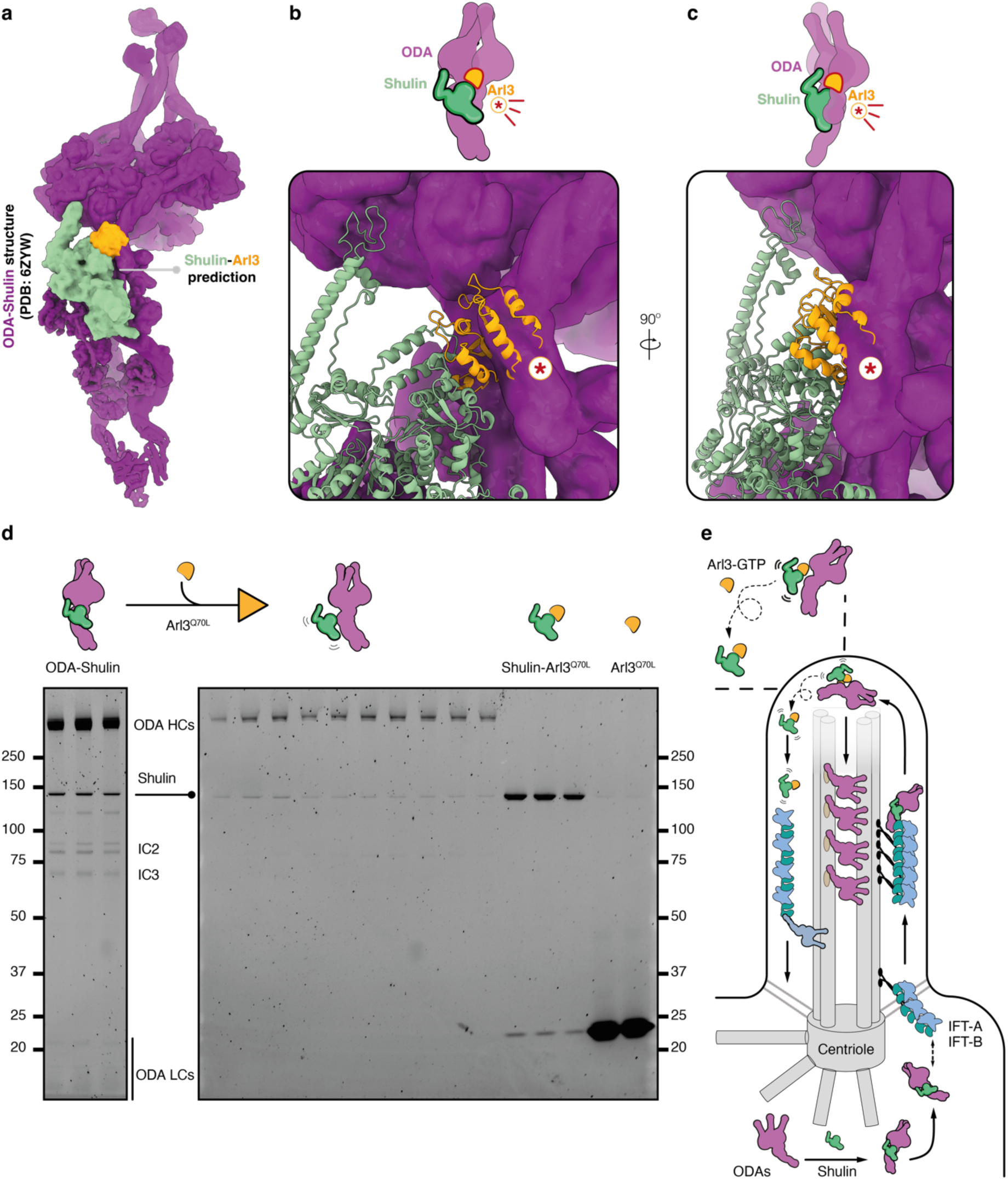
Active Arl3 displaces Shulin from ODAs. **a.** Shulin-Arl3 prediction generated using AF3 superimposed on the experimental structure of ODA-Shulin complex (PDB: 6ZYW). **b, c**. Structural models and cartoon representations highlighting steric clashes (encircled asterisk) between Arl3 and ODAs that would prevent formation of a stable ODA-Shulin complex. **d.** Assay to test displacement of Shulin from a purified ODA-Shulin complex by adding excess of the GTP mimetic Arl3^Q70L^ variant is shown as a schematic above gel images. The left-hand gel image shows size-exclusion chromatography fractions from the peak containing the purified ODA-Shulin complex. The right-hand gel image shows size-exclusion chromatography fractions following injection of the purified ODA-Shulin complex incubated with excess Arl3^Q70L^. **e.** Proposed model highlighting Shulin’s role in delivering ODAs to motile cilia. Shulin (DNAAF9) locks ODAs in a closed conformation and chaperones their attachment to IFT trains near the basal body for ciliary entry. Inside cilia, Shulin (DNAAF9) is displaced by active Arl3-GTP for ODA activation and stable axonemal incorporation. Shulin (DNAAF9) could be sequestered in a complex with Arl3 that exits cilia for recycling or degradation in the cytoplasm.

## Discussion

Our work provides a molecular explanation of how ODAs get activated in cilia. We show that the packaging chaperone Shulin is displaced from *Tetrahymena* ODAs by the ciliary GTPase Arl3 to promote motor activation. Based on the conservation of Arl3’s interaction with Shulin and its human orthologue DNAAF9, we propose that the mechanism of ODA activation is conserved in several ciliates.

We determine a key function of DNAAF9 in ODA transport by showing it interacts with intraflagellar transport proteins in human airway cells. Based on its interactions and the sub-cellular concentration of DNAAF9 at the base of airway cilia, we speculate that it could aid the attachment of packaged ODAs to IFT trains near the basal body region for ciliary entry. The precise mechanism of how ODAs bind IFT trains remains poorly resolved and will be the focus of future work.

DNAAF9’s interaction with ARL3, a ubiquitously expressed GTPase, suggests that in metazoans, DNAAF9 could be important for functions beyond the one’s we define here. The DNAAF9-ARL3 interaction was first detected in retinal RPE-1 cells that have primary cilia. DNAAF9’s broad expression pattern in humans also indicates it could play roles in tissues with both primary and motile cilia. Further studies are needed to address its importance to cilia functions more broadly. DNAAF9 variants have not been found in PCD patient cohorts so far and it is possible that its loss severely impacts the functions of several cilia sub-types. Therefore, we suggest that DNAAF9 should be considered as a novel candidate for syndromic ciliopathies with combined features characteristic of primary and motile cilia defects.

We extend ARL3’s functions to include the regulation of ODA transport and their timely activation in motile cilia. Complete loss of ARL3 is lethal in vertebrates^24^ and loss of function ARL3 variants have been classically linked to severe primary ciliopathies due to perturbed trafficking of ciliary signalling receptors^12, 25, 26^. Arl3 is essential for the formation of motile flagella in *Leishmania*^27^. Recent work using *T. brucei* has also identified an interaction between Arl3 and the ODA transport adaptor Oda16 where the latter is proposed to act as an effector for the ciliary transport of motility related cargoes including ODAs and the central pair component HYDIN^28^. Taken together, our work and these other findings indicate that there is a complex interplay between ARL3 and ODA transport factors such as DNAAF9/Shulin and Oda16 during motile cilia formation which should be explored in future work. Given these emerging links between ARL3 and the biogenesis of motile cilia, we recommend that primary ciliopathy patients carrying ARL3 variants should be additionally assessed for clinical features indicative of defective motile cilia.

The sheer scale of axonemal dynein biosynthesis and transport in multiciliated cells could necessitate multiple adaptors functioning over several steps to concentrate ODAs around and inside growing cilia. Crucially, the need to shut off ODA motor activity is key to preventing aberrant off-target interactions along the trafficking route, highlighting the critical inhibitory role played by Shulin (DNAAF9). Once inside cilia, we propose that Arl3 in its GTP bound conformation promotes the release of Shulin (DNAAF9) from ODAs and sequesters it to prevent re-binding (**Fig. 5e**). The DNAAF9-ARL3 complex could diffuse out of cilia for recycling or degradation. Our displacement model is compatible with ODAs undergoing an irreversible structural reconfiguration from a ‘closed’ (inhibited) to an ‘open’ (active) state specifically inside cilia during their stable incorporation into a growing ciliary axoneme.

## Supporting information

Supplementary Raw Data File 1_Arl3DisplacesShulinToActivateCiliaryODAs

Supplementary Raw Data File 2_Arl3DisplacesShulinToActivateCiliaryODAs

## Extended data figures and legends

**FIGURE S1.**
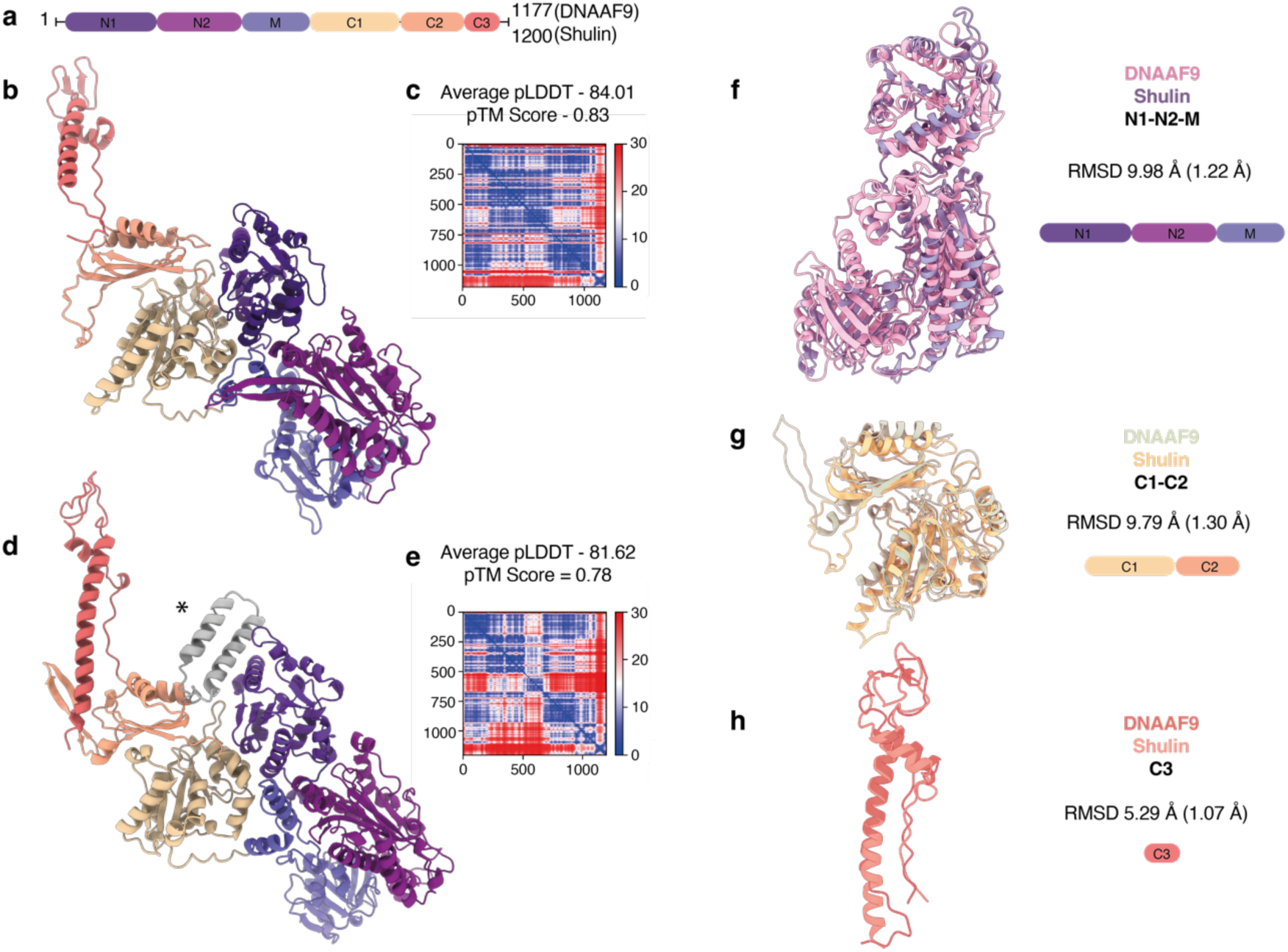
DNAAF9 and Shulin share high overall and domain level structural similarity. **a.** Domain organisation of DNAAF9 and Shulin. **b, c, d, e.** AlphaFold2 generated models with PAE plot, pLDDT and pTM scores for DNAAF9 and Shulin. **f, g, h.** Structural comparison of DNAAF9 and Shulin divided into sub-domains (f: N1-N2-M domains, g: C1-C2 domains, h: C3 extension) by RMSD analysis. Overall RMSD values are shown with scores for best aligned regions included in brackets. Flexible loop regions were retained for the analysis. * An extra helix-loop-helix (gray) is predicted in Shulin’s structure in (d); this was excluded for calculating domain level RMSD scores.

**FIGURE S2.**
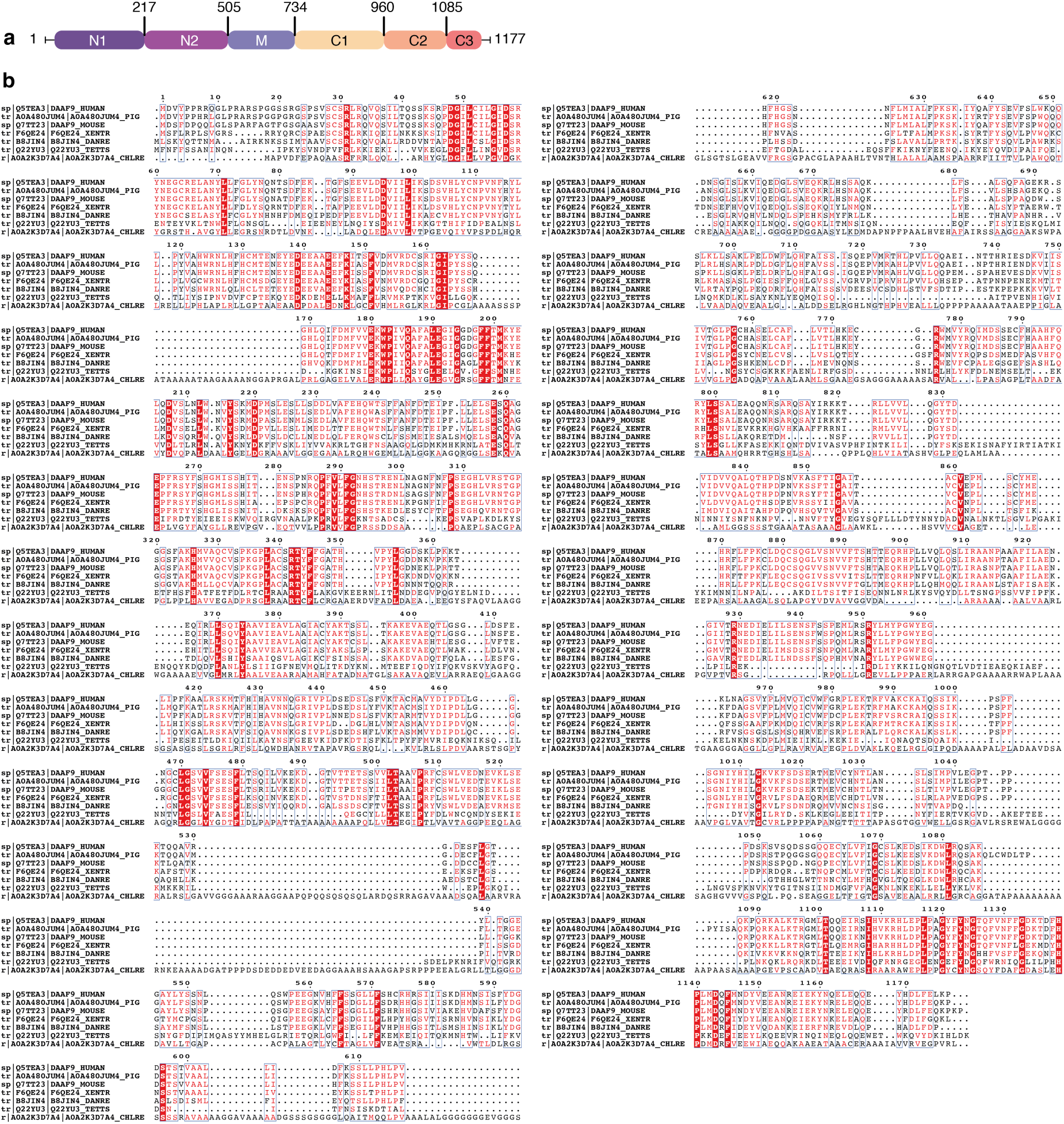
DNAAF9 domain architecture and multiple sequence alignment with its orthologs. **a.** Domain boundaries of DNAAF9 **b.** Sequence alignment between DNAAF9 and its orthologs across model ciliate species generated using ESPript. UniProt IDs are labelled to the left. Conserved residues are highlighted in red and similar residues are colored in red with blue boxes around.

**FIGURE S3.**
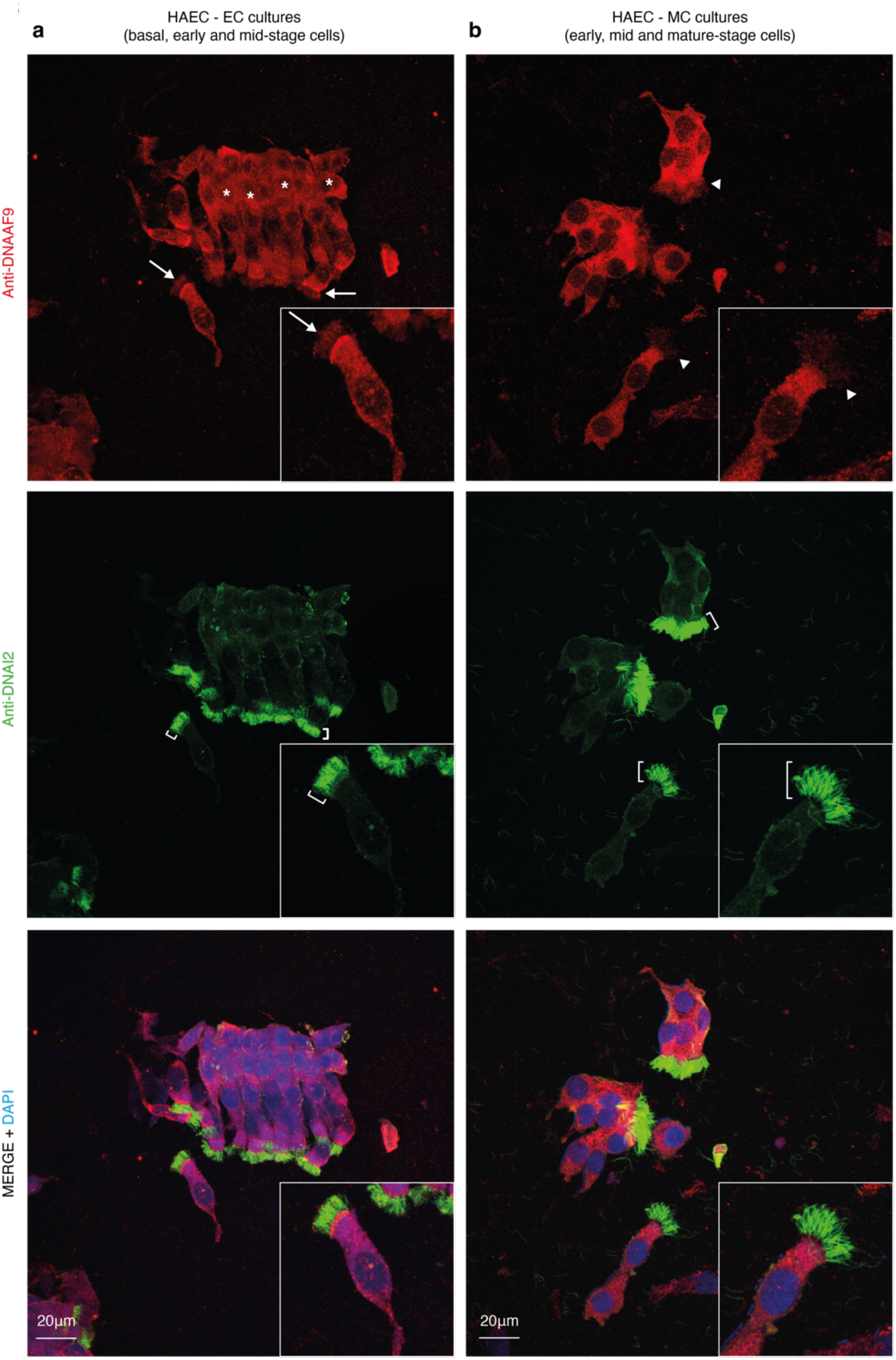
Co-immunostaining of DNAAF9 and DNAI2 in ECs and MCs. **a., b.** Co-immunostaining of DNAAF9 and DNAI2 in ECs with short cilia (arrowhead and short brackets) and MCs with long cilia (white arrowhead and long brackets). Asterisks * in (a) show non-ciliated basal cells in ECs. Inset panels show an enlarged image of an EC and MC for comparison.

**FIGURE S4.**
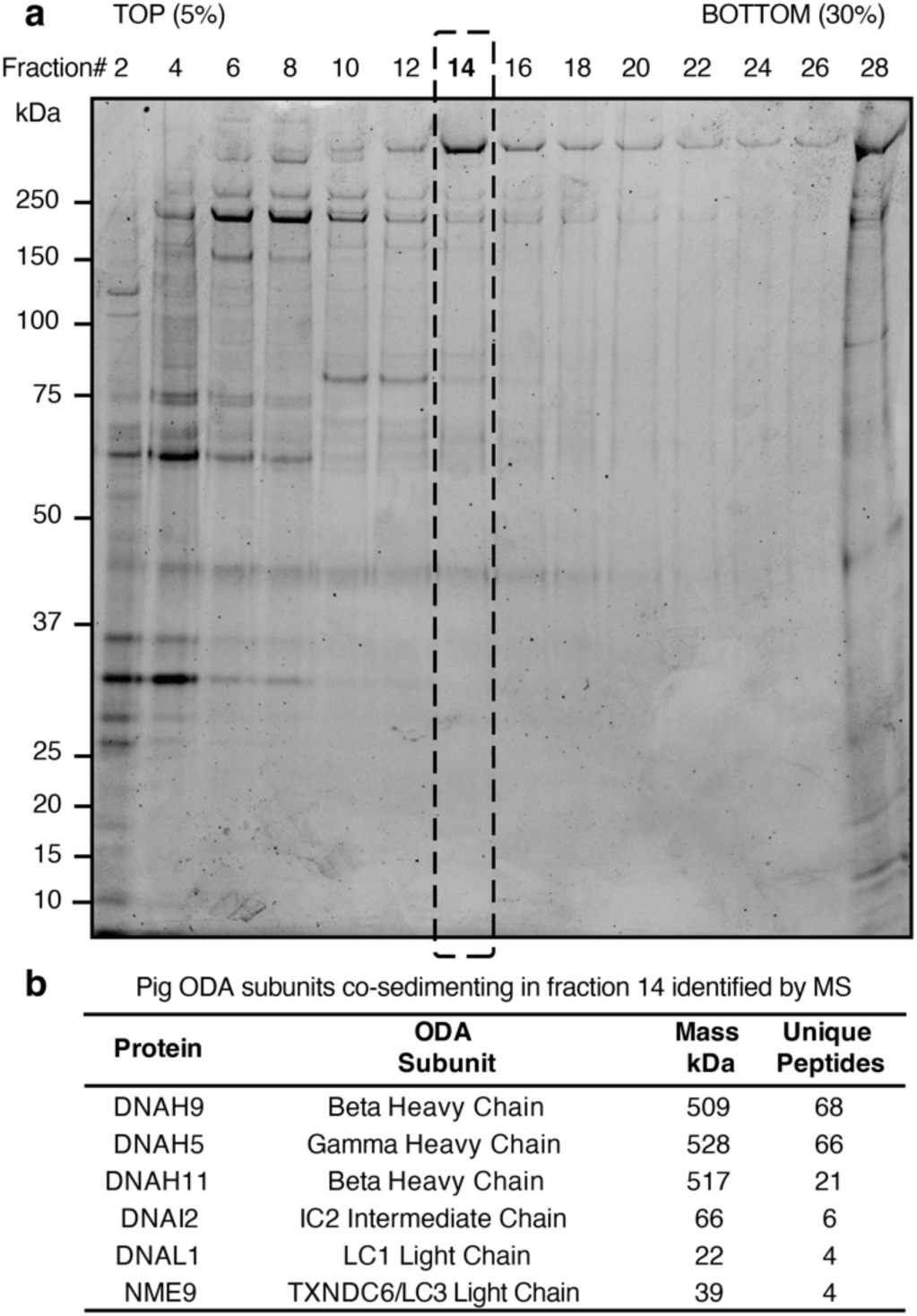
Purification of ODAs from pig tracheal cilia. **a.** SDS-PAGE analysis of fractions from a sucrose density gradient centrifugation resolving complexes obtained from high-salt extraction of pig tracheal cilia axonemes. Fraction 14 shows a major high molecular weight band consistent with the mass of a dynein heavy chain. **b.** Mass spectrometry analysis of ODA subunits co-sedimenting in fraction 14. Unique peptide counts detected at a 95% peptide threshold are shown.

**FIGURE S5.**
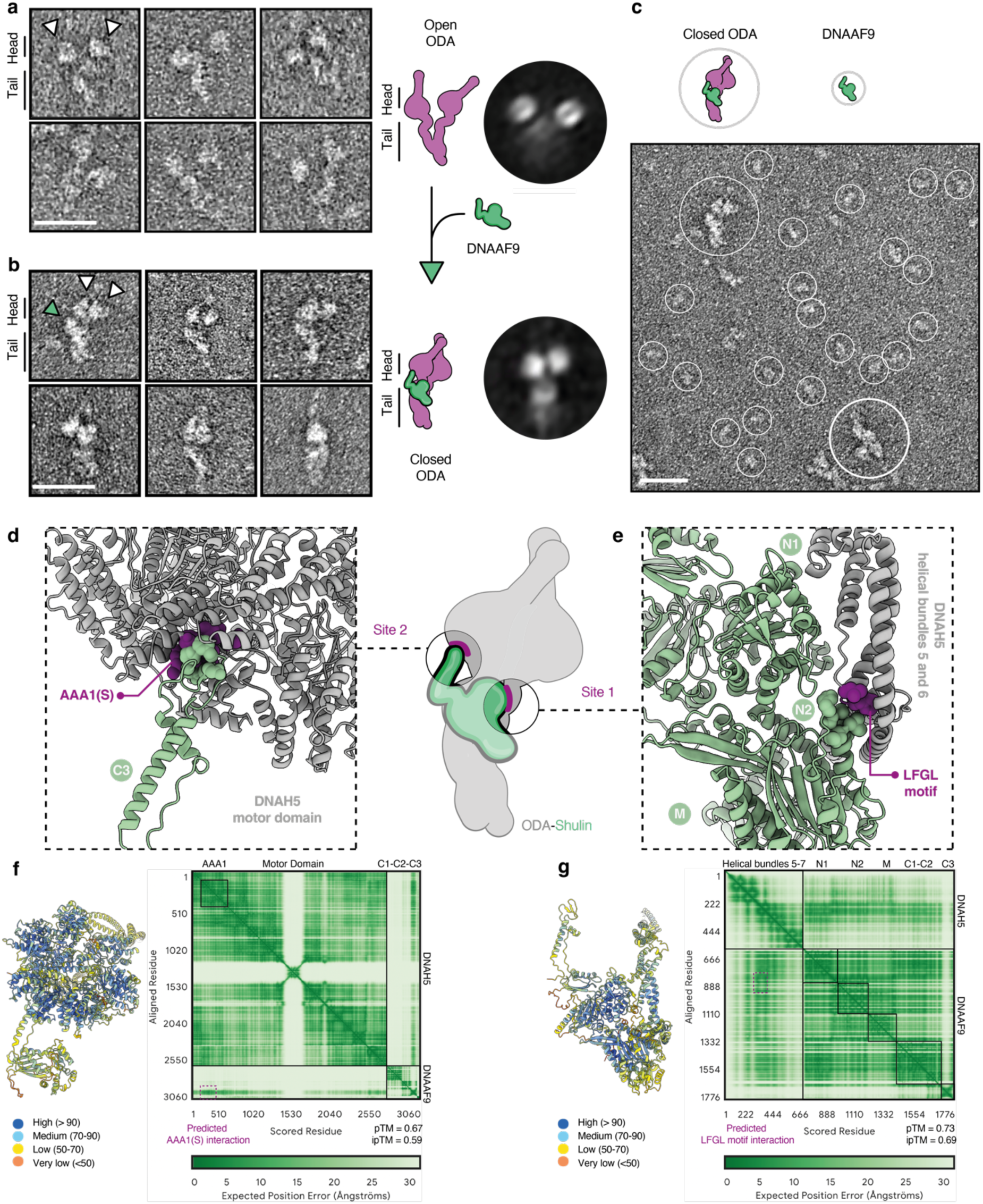
DNAAF9 binds mammalian ODAs to enforce conformational closure. **a, b.** Electron micrographs of negatively stained two-headed pig ODA particles in isolation and when combined with recombinant DNAAF9. Particles adopting an open conformation are shown in (a) and those with a closed phi-particle like conformation are in (b) with white arrowheads marking the positions of the motor heads in top left panels; the green arrowhead in top left panel in (b) points to a putative density for DNAAF9. Scale bars, 50nm. Conformational differences between the two samples are illustrated by a schematic diagram and representative 2D class averages highlighting the open and closed states are shown to the right. **c.** A representative negative stain electron micrograph with single particles of pig ODA holocomplexes and DNAAF9 encircled. Scale bar, 25nm. **d, e.** AlphaFold3-Multimer model of key binding interfaces (site 1 and 2) between DNAAF9 and DNAH5 (gamma heavy chain of human ODAs). Conserved interface residues at the AAA1 site in the motor domain and the LFGL motif in the tail helical bundle of DNAH5 are surface represented as purple spheres with the corresponding contacting residues in DNAAF9 shown as green spheres. **f, g.** pLDDT scores are plotted onto the predicted structures to highlight local confidence in the modeling (dark blue = pLDDT 90-100; high confidence; orange = pLDDT 0-50; low confidence). PAE plots highlighting interaction interfaces and model statistics are shown.

**FIGURE S6.**
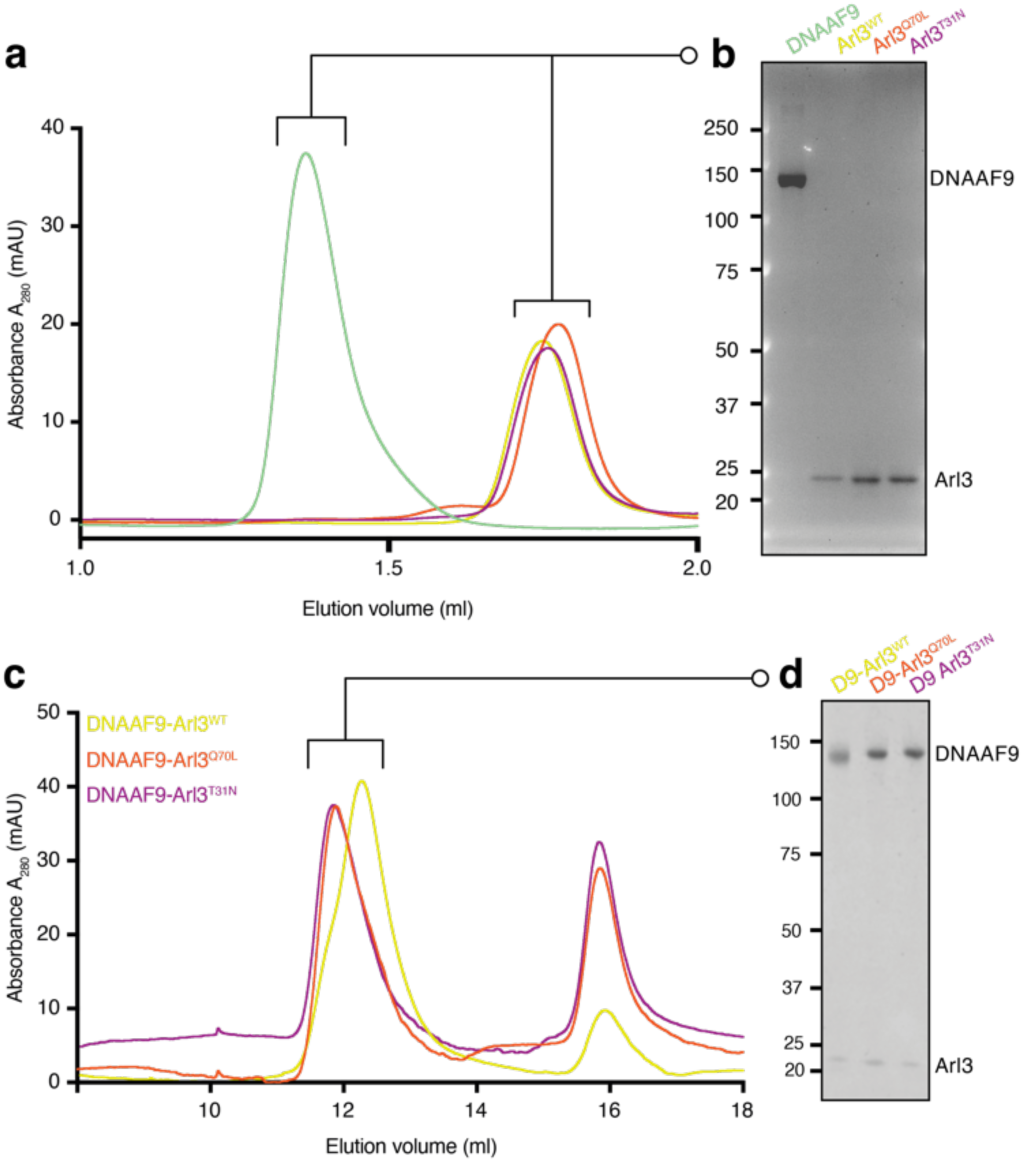
Purification of recombinant DNAAF9 and Arl3 variants and complex reconstitutions in the presence of GTP. **a, b.** Size exclusion chromatography profiles of recombinant DNAAF9 and Arl3 variants expressed and purified from insect cells over a Superose 6 10/300 column are shown. Gel image to the right shows SDS-PAGE analysis of peak fractions. **c. d.** Reconstitutions of DNAAF9 with three Arl3 variants [wildtype, GTP (Q70L) and GDP (T30N) mimicking mutants] in 1mM GTP. SDS-PAGE analysis shows proteins co-eluting in the first peak in a complex from each of the three reconstitutions.

**FIGURE S7.**
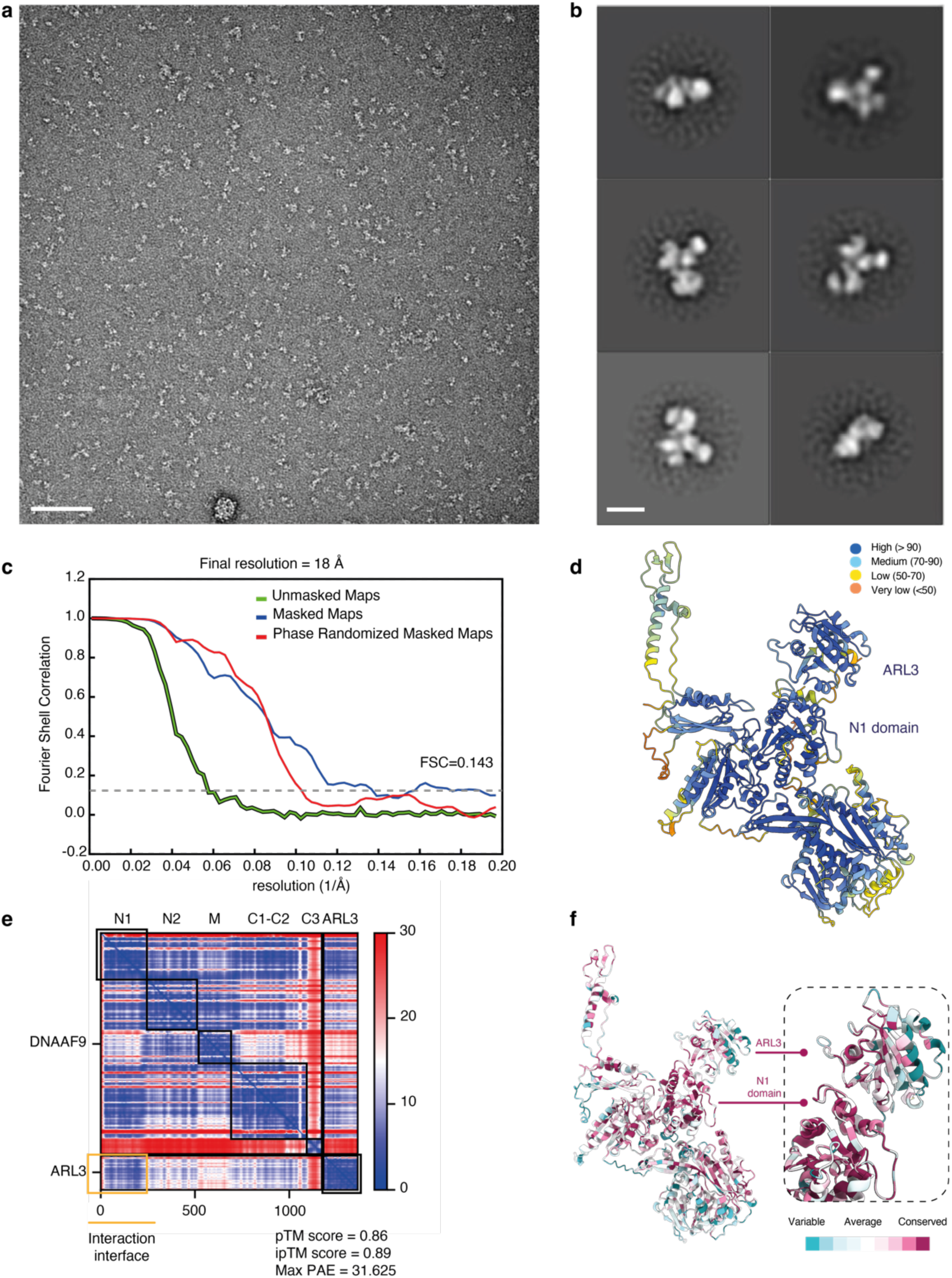
Negative stain EM, AlphaFold2-Multimer and CONSURF analysis of the DNAAF9-ARL3 complex. **a.** A representative negative stain electron micrograph showing single particles from a DNAAF9-Arl3^Q70L^ reconstitution experiment. Scale bar, 100 nm. **b.** Representative class averages of DNAAF9 bound by Arl3^Q70L^ from 2D classification of the reconstituted complex. Scale bar, 2.5 nm **c.** Gold standard Fourier shell correlation (FSC) curves for DNAAF9-ARL3 complex map (Fig. 4b) as determined by RELION-4.0 (FSC=0.143). **d.** pLDDT scores are plotted onto the predicted DNAAF9-ARL3 structure to highlight local confidence in the modeling (dark blue = pLDDT 90-100; high confidence; orange = pLDDT 0-50; low confidence). **e.** PAE plot from AF2-Multimer prediction of the DNAAF9-ARL3 complex. The interaction interface is highlighted, and model statistics are shown. **f.** CONSURF scores mapped onto the predicted model of the DNAAF9-ARL3 complex.

**FIGURE S8.**
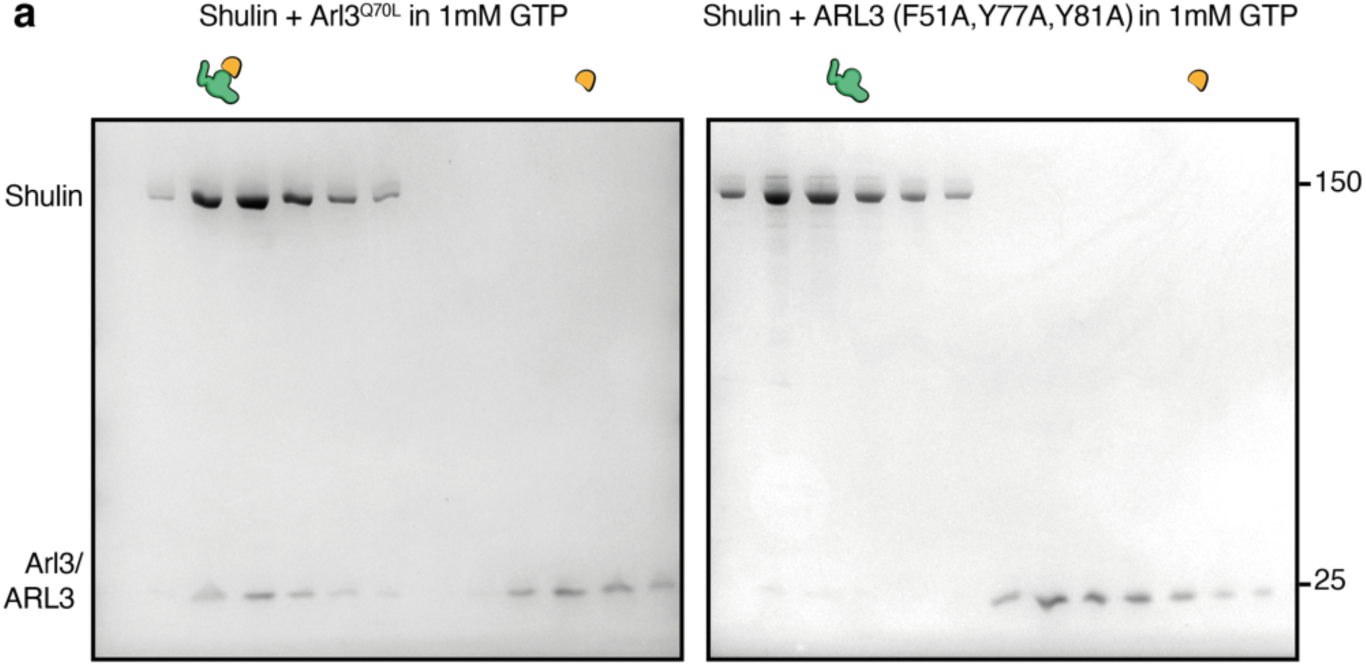
Mutating the interaction interface destabilises Shulin-ARL3 complex. **a.** Analytical size-exclusion chromatography fractions from characterisations of complex formation between Shulin and wildtype Arl3 or the triple alanine substitution ARL3 mutant.

**FIGURE S9.**
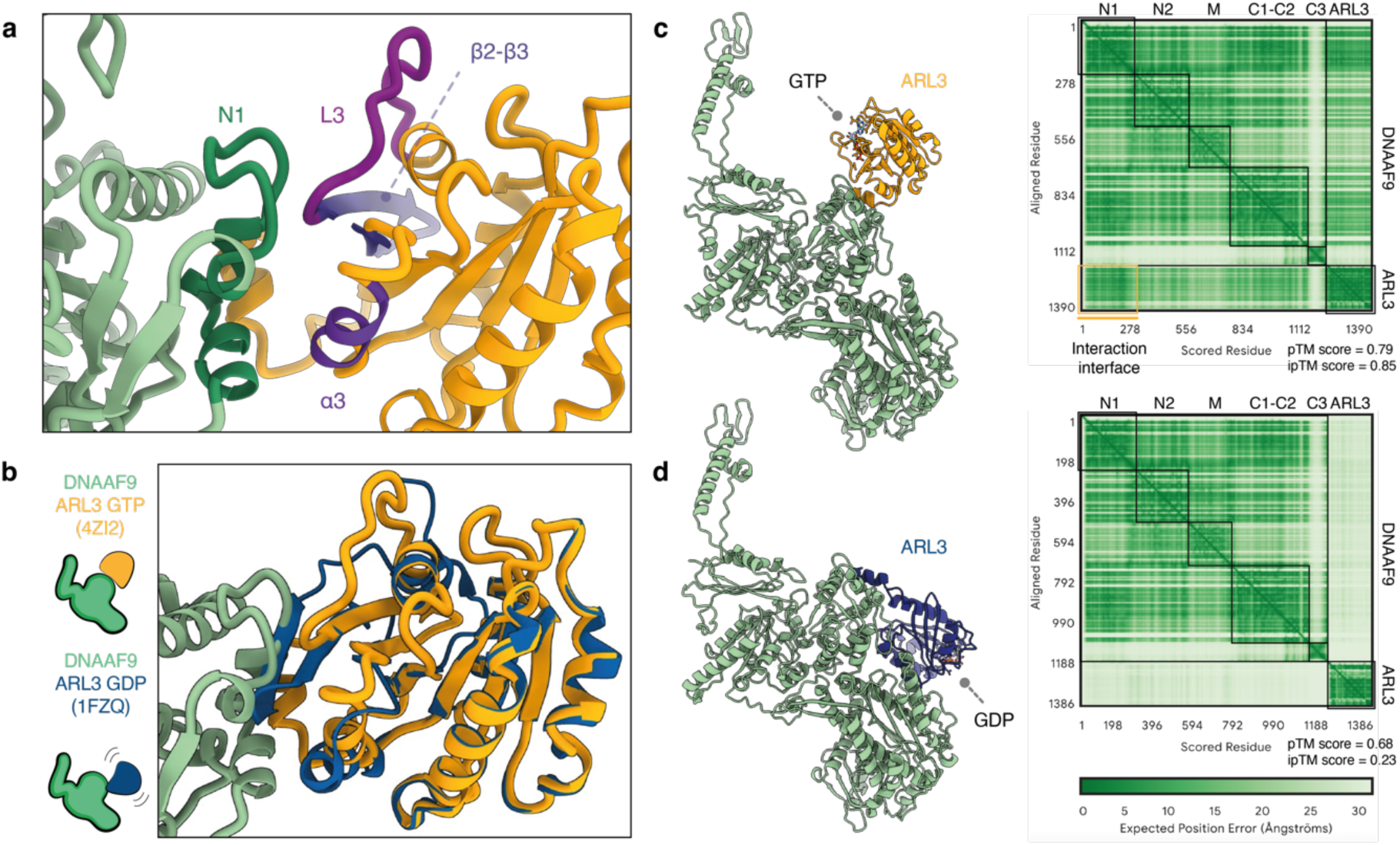
Steric clashes between ARL3-GDP and DNAAF9 are incompatible for complex formation. **a.** Secondary structure features in ARL3’s GTP loaded state that interact with DNAAF9’s N1 domain are shown. **b.** Structure of ARL3’s in its GDP loaded state (blue) superimposed on the AF2-Multimer prediction of the DNAAF9-ARL3 complex. ARL3’s L3 loop and β2 strand adopts a beta-hairpin in the GDP state causing steric clashes with DNAAF9. **c. d.** AF3-Multimer models of DNAAF9-ARL3 in the presence of GTP and GDP ligands. PAE plots are shown to the right of each prediction. The interaction interface is highlighted, and model statistics are shown.

## Materials and Methods

### Immunostaining and image analysis

Human airway epithelial cell (HAEC) cultures were purchased from Epithelix Sarl at day 17 post airlift (early to mid) and day >45 post airlift (mid to mature). Cells were spread on superfrost slides, air-dried, and processed for immunofluorescence. HAECs were fixed for 15 minutes with 4% PFA and then permeabilised with 0.25% PBST (Triton-X 100) for 10 minutes permeabilization. Cells were blocked in 10% donkey serum in PBST for one hour at room temperature and incubated with primary antibodies against DNAAF9 (Proteintech, 23184-1-AP), DNAI2 (H00064446-M01, clone IC8) in PBST and 1% donkey serum either for one hour at room temperature or at 4’C overnight. Cells were incubated in secondary antibody for 1 hour following three rounds of 15-minute PBST washes. After final three 15-minute washes in PBST cells were counterstained and mounted under a 1.5mm glass coverslip using prolong gold antifade medium with DAPI. Corners of the coverslip were sealed with commercial nail varnish. All steps were performed at room temperature.

Cells were imaged with a Leica SP8 Confocal Laser Scanning Microscope with a 63x HC PL APO CS2 immersion oil objective. Fluorophores were excited at 488nm and 568nm using a 50mW 405nm diode and 65mW argon laser. Lasers were set at 2-5% intensity with an acquisition pinhole of 1. Number of pixels per image was set by first imaging the larger more elongated late to mature stage cells and used for smaller and rounder/less elongated early to mid-stage cells. All images were acquired at room temperature using the same parameters for image analysis and quantification.

Image analysis was performed using Fiji ImageJ^29^. Ciliated cells that had no overlapping regions with other cells were included in the analysis. Cilia length measurements were performed by first defining cilia regions as the structure spanning from the first visible apical DNAI2 signal to the ciliary tips. Splayed cilia that appeared detached from the cell body were excluded. Using the ImageJ measuring tool, 3 distinct cilia regions were measured across the apical length to obtain the average cilia length. Signal intensity measurements were performed by demarcating cilia only, cell only and background regions using the Freehand Region of Interest (ROI) tool to obtain a cilia, cell, and background ROI. Measurements were set to analyse area, integrated density and mean gray values, which were used to calculate the corrected total cilia fluorescence (CTCF) where CTCF = Integrated density – (cilia area x mean background fluorescence). The mean background fluorescence used above was calculated by obtaining background intensity with cell bodies included in the ROI without the cilia ROIs and then subtracting with background intensity excluding the cell body and the cilia ROIs. CTCF values were Log2 transformed and plotted in GraphPad Prism (LaJolla, CA) software.

### Immunoprecipitation, mass spectrometry and immunoblotting

Endogenous DNAAF9 immunoprecipitations were performed using HAECs (MucilAir, Epithelix Sarl) at early to mid (17 days post-ALI) and mid to mature (40 days post-ALI) stages of differentiation. Whole cell lysates were obtained by incubating cells in IP lysis buffer (50 mM Tris-HCl pH 7.5, 100 mM NaCl, 10% Glycerol, 0.5 mM EDTA, 0.5% IGEPAL. An EDTA free protease inhibitor tablet, 5uM MG132 and 1mM DTT. Cells were passed through a 25-gauge needle several times to ensure efficient homogenisation and lysis. For preserving potential HSP90 interactions, 25mM sodium molybdate (Sigma-Aldrich) was included in the IP lysis buffer to reduce ATP hydrolysis and client release. Equivalent concentration of whole cell lysates was obtained from early-mid and mid-mature stage cultures (13.7 mg/ml and 14.0 mg/ml respectively). Lysates from both cell stages were incubated for up to 4 hours at 4°C with an antibody against DNAAF9/C20ORF194 (Proteintech, 23184-1-AP) and an isotype-matched IgG rabbit polyclonal antibody (Proteintech, 30000-0-AP) as control. Immunocomplexes in the antibody-lysate solutions were concentrated onto Pierce Protein A/G magnetic beads (Thermo Fisher) by incubating for 30 minutes at 4°C. Two washes were performed in the IP lysis buffer followed by two washes in the same buffer but with the IGEPAL concentration reduced to 0.2%. Final two washes were performed in the IP lysis buffer without any detergent.

Following washes, beads were frozen at −20°C for subsequent on-bead tryptic digestion and mass-spectrometric analysis. For immunoblotting analyses, immunocomplexes were eluted by boiling the beads in elution buffer (20% LDS with 10% DTT in Milli Q water) and resolved on NuPAGE Novex 4– 12% Bis–Tris Protein Gels (Life Technologies). Proteins were transferred onto PVDF membranes using the XCell II Blot module (Life Technologies) and blocked in 5% BSA followed by immunoblotting with the following antibodies: anti-DNAAF9 (Proteintech, 23184-1-AP), anti-DNAI2 (M01, clone IC8; Abnova), anti-DNAH5 (Sigma, HPA037470), anti-IFT74 (Proteintech, 27334-1-AP), anti-IFT81 (Proteintech, 11744-1-AP) and anti-DYNC1I2 (Proteintech, 12219-1-AP). All primary antibody dilutions were in the range of 1/500 – 1/1000 in PBST (0.1% Tween-20) with 1% BSA. Membranes were washed three times in PBST (0.1% Tween-20) and incubated with HRP conjugated rabbit or mouse secondary antibodies (GeneTex, EasyBlot anti-rabbit IgG and anti-mouse IgG HRP conjugated monoclonal antibodies; GTX221666-01-S and GTX221667-01-S respectively) at 1/1000 dilution in PBST (0.1% Tween-20) with 1% BSA. Protein bands were detected with SuperSignal West Femto reagent (Thermo Scientific) and membranes were visualised on an Odyssey Fc Imager (LI-COR Biosciences) or an iBright FL1000 Imaging System under chemiluminescence settings (Thermo Fisher).

### Proteomics

#### TMT Labelling and High pH reversed-phase chromatography

Immunoprecipitated protein samples were reduced (10mM TCEP, 55°C for 1h), alkylated (18.75mM iodoacetamide, room temperature for 30min.) and then digested from the beads with trypsin (1.25µg trypsin; 37°C, overnight). The resulting peptides were then labeled with Tandem Mass Tag (TMTpro) sixteen plex reagents according to the manufacturer’s protocol (Thermo Fisher Scientific, Loughborough, LE11 5RG, UK) and the labelled samples pooled and desalted using a SepPak cartridge according to the manufacturer’s instructions (Waters, Milford, Massachusetts, USA). Eluate from the SepPak cartridge was evaporated to dryness and resuspended in buffer A (20 mM ammonium hydroxide, pH 10) prior to fractionation by high pH reversed-phase chromatography using an Ultimate 3000 liquid chromatography system (Thermo Scientific). In brief, the sample was loaded onto an XBridge BEH C18 Column (130Å, 3.5 µm, 2.1 mm X 150 mm, Waters, UK) in buffer A and peptides eluted with an increasing gradient of buffer B (20 mM Ammonium Hydroxide in acetonitrile, pH 10) from 0-95% over 60 minutes. The resulting fractions (6 in total) were evaporated to dryness and resuspended in 1% formic acid prior to analysis by nano-LC MSMS using an Orbitrap Fusion Lumos mass spectrometer (Thermo Scientific).

#### Nano-LC Mass Spectrometry

High pH RP fractions were further fractionated using an Ultimate 3000 nano-LC system in line with an Orbitrap Fusion Lumos mass spectrometer (Thermo Scientific). In brief, peptides in 1% (vol/vol) formic acid were injected onto an Acclaim PepMap C18 nano-trap column (Thermo Scientific). After washing with 0.5% (vol/vol) acetonitrile 0.1% (vol/vol) formic acid peptides were resolved on a 250 mm × 75 μm Acclaim PepMap C18 reverse phase analytical column (Thermo Scientific) over a 150 min organic gradient, using 7 gradient segments (1-6% solvent B over 1min., 6-15% B over 58min., 15-32%B over 58min., 32-40%B over 5min., 40-90%B over 1min., held at 90%B for 6min and then reduced to 1%B over 1min.) with a flow rate of 300 nl min^−1^. Solvent A was 0.1% formic acid and Solvent B was aqueous 80% acetonitrile in 0.1% formic acid. Peptides were ionized by nano-electrospray ionization at 2.0kV using a stainless-steel emitter with an internal diameter of 30 μm (Thermo Scientific) and a capillary temperature of 300°C. All spectra were acquired using an Orbitrap Fusion Lumos mass spectrometer controlled by Xcalibur 3.0 software (Thermo Scientific) and operated in data-dependent acquisition mode using an SPS-MS3 workflow. FTMS1 spectra were collected at a resolution of 120 000, with an automatic gain control (AGC) target of 200 000 and a max injection time of 50ms. Precursors were filtered with an intensity threshold of 5000, according to charge state (to include charge states 2-7) and with monoisotopic peak determination set to Peptide. Previously interrogated precursors were excluded using a dynamic window (60s +/-10ppm). The MS2 precursors were isolated with a quadrupole isolation window of 0.7m/z. ITMS2 spectra were collected with an AGC target of 10 000, max injection time of 70ms and CID collision energy of 35%. For FTMS3 analysis, the Orbitrap was operated at 50,000 resolutions with an AGC target of 50,000 and a max injection time of 105ms. Precursors were fragmented by high energy collision dissociation (HCD) at a normalised collision energy of 60% to ensure maximal TMT reporter ion yield. Synchronous Precursor Selection (SPS) was enabled to include up to 10 MS2 fragment ions in the FTMS3 scan.

#### Data Analysis

The raw data files were processed and quantified using Proteome Discoverer software v2.4 (Thermo Scientific) and searched against the UniProt Human database (downloaded January 2023: 81579 entries) using the SEQUEST HT algorithm. Peptide precursor mass tolerance was set at 10ppm, and MS/MS tolerance was set at 0.6Da. Search criteria included oxidation of methionine (+15.995Da), acetylation of the protein N-terminus (+42.011Da) and Methionine loss plus acetylation of the protein N-terminus (−89.03Da) as variable modifications and carbamidomethylation of cysteine (+57.0214) and the addition of the TMTpro mass tag (+304.207) to peptide N-termini and lysine as fixed modifications. Searches were performed with full tryptic digestion and a maximum of 2 missed cleavages were allowed. The reverse database search option was enabled, and all data was filtered to satisfy false discovery rate (FDR) of 5%.

#### Proteomics data processing

Mass spectrometry datasets were processed in MS Excel. Protein abundances for each replicate run were used to rank proteins by log_2_ fold-change (enrichment) between DNAAF9 IPs over IgG controls. A two-tailed, unpaired t-test assuming equal variances was performed to obtain a p-value statistic for each protein from triplicate datasets per differentiation stage to rank the hits by significance of enrichment in DNAAF9 IPs versus IgG controls (**Supplementary raw data file 1**). Proteins were filtered against the CRAPome repository set for human proteins (http://www.crapome.org/,^30^) using an arbitrary threshold of 50 i.e. proteins appearing in >50 out of 716 proteomics experiments using human samples captured in CRAPome. All protein hits with CRAPome values higher than 50 were removed from further analyses except those that had fold-change enrichment of >2.5 (log_2_ fold change of 1.32) and p-values > 0.05 (-log10 p-value 1.30) in the DNAAF9 IPs. These protein hits were included in further analyses and taken forward for validation to account for high abundance ‘true’ interactors. DNAAF9 interactomes from immature and mature cell stages were visualised as scatter plots (log2 fold-change versus -log10 p-value) using GraphPad Prism9.

Three types of data analyses and annotation were performed using the proteomics datasets. First, a comparison of the early and late stage interactomes was performed to curate a list of proteins that interacted with DNAAF9 at both stages of differentiation. Gene Ontology analysis on this list of common 87 protein hits was performed using ShinyGO to represent enrichment against a background list of all human protein-coding genes based on cellular component (GO:0005575)^31^. Second, the abundances between the most highly enriched interactors were compared between early and mid-stages using ProHits-viz^32^ and fold changes, changes in relative abundances and significance for each protein interactor were visualised. Finally, unique protein interactors that were found exclusively in either the early or late-stage datasets were identified, and bioinformatics searches were performed to annotate their cellular localisations.

### Identification of proteins by mass spectrometry

Two separate types of MS analyses were performed for protein identification. For identifying interactors of the GTP locked ARL3^Q70L^ variant in *Tetrahymena*, eluates from strep pull-downs were resolved on an SDS-PAGE gel and silver stained to visualise bands for co-eluting proteins. In parallel, the same eluates were tryptically digested in-solution for mass spectrometry analysis (**Supplementary raw data file 2**). For identifying pig ODA subunits co-fractionating together in fraction 14 of the 5-30% sucrose density gradient, sypro stained gel slices were excised from the entire lane corresponding to fraction 14 and subjected to in-gel tryptic digestion followed by mass spectrometry analysis (**Supplementary raw data file 2**).

### Cloning and protein expression in insect cells or bacteria

All genes were codon-optimized for insect cell or bacterial expression. Gene sequence coding for *Tetrahymena thermophila* Shulin (Q22YU3) was expressed in insect cells using Addgene plasmid #170315 and purified as described previously (Mali et al., 2021). *Tetrahymena thermophila* small GTPase Arl3 (Q229S0) codon optimised gene sequences coding for the wildtype and the Q70L and T30N variants and human C20orf194/DNAAF9 (Q5TEA3) were gene synthesized (Epoch Life Science Inc.) and cloned in pACEBac1 vectors containing a C-terminal 2xStrep tag using Gibson assembly for insect cell expression. Human ARL3 (P36405) triple mutant (ARL3 F51A-Y77A-Y81A) was gene synthesized and cloned into a pET22b (+) vector containing a C-terminal His tag for periplasmic bacterial expression (Epoch Life Science Inc.).

For insect cell expression, plasmids were first amplified by transforming chemically competent DH5α cells (Thermo Scientific). To generate baculovirus DNA (bacmid), chemically competent DH10 EmBacY cells (Thermo Scientific) were transformed and recombinant bacmids were selected by blue/white screening. Purified bacmids carrying the gene of interest were used to transfect *Spodoptera frugiperda* derived Sf9 cells with FuGENE following the supplier’s instructions (Promega). At day 4 post-transfection, infected Sf9 cells were inspected under a fluorescence microscope for YFP expression (yellow fluorescence protein) to monitor virus production. Low-titer P1 baculoviruses were collected from the culture suspension and used for virus amplification to produce P2 baculoviruses at day 7 post-transfection. For recombinant protein production, 5 mL of P2 baculoviruses were used to inoculate 500 mL of Sf9 cells at a density of 2 x 10^6^/mL in *SF 900 II SFM* media (Gibco). Cells were harvested at day 3 post-infection by centrifuging at 2,000xg for 15 mins at 4°C using a JLA 8.1000 rotor (Beckman Coulter). Insect cell pellets were frozen in liquid nitrogen for storage or used immediately for protein purification.

For bacterial cell expression, the plasmid pET22b (+) carrying ARL3-F51A-Y77A-Y81A (ARL3-FYY) was transformed into chemically competent BL21 (DE3) cells (Thermo Scientific). A saturated overnight pre-culture of BL21 (DE3) cells (OD_600nm_ 0.05) was used to inoculate 4 liters of Luria Broth (LB) media (Sigma) supplemented with 100 μg/mL of Ampicillin (Sigma) at 37°C. At an OD_600nm_ between 0.6 – 0.8, expression was induced by adding isopropyl β-D-thio-galactopyranoside (IPTG) at a final concentration of 0.5 mM. Upon induction, the temperature was decreased to 18°C for overnight protein expression. Cells were harvested by centrifuging at 5,000xg for 15 mins at 4°C using a JLA 8.1000 rotor (Beckman Coulter). Bacterial cell pellets were frozen in liquid nitrogen for storage or used immediately for protein purification.

### Protein purification

Human C20orf194/DNAAF9 (Q5TEA3), *Tetrahymena thermophila* Shulin (Q22YU3) and Arl3 (Q229S0) proteins [WT, Q70L, T30N] were purified from *Sf9* cells as described previously (Mali et al., 2021). Briefly, cell pellets were mechanically lysed in 20 mL lysis buffer (20 mM Hepes-NaOH (pH 7.2), 100 mM NaCl, 2 mM MgAc, 1 mM EDTA, 10% (v/v) glycerol, 1 mM DTT) using a Dounce homogenizer (Wheaton) for up to 100 strokes on ice. Lysates were clarified by ultracentrifugation at 47,000xg for 45 mins, at 4°C in a Ti70 rotor (Beckman Coulter). Clarified lysates were loaded using a sample pump or super-loop (Cytiva) on a 5 mL Strep-Trap HP column (GE Healthcare) which was pre-equilibrated with 5 column volumes (CV) of lysis buffer. The resin was washed with 10 CV of lysis buffer to remove non-specific binding of contaminants. Recombinant proteins were eluted off the resin by passing 5 CV of the elution buffer (lysis buffer containing 3 mM Desthiobiotin (IBA). Eluates were resolved on NuPAGE Novex 4–12% Bis–Tris Protein Gels (Life Technologies). and stained with Sypro ruby (Invitrogen) to assess the purity of the recombinant proteins. Eluted proteins were further purified over gel filtration at 4°C on a Superdex 200 10/300 GL (GE Healthcare) column the using lysis buffer. Purified proteins eluted in single peaks and were snap frozen in liquid nitrogen for storage at −70°C.

His-tagged human ARL3 (P36405) triple mutant was purified from bacterial pellets. Pellets were resuspended in 35 mL lysis buffer (50 mM Hepes pH 7.4, 200 mM NaCl, 5% Glycerol, 1 mM MgCl2, 1 mM DTT) and lysed on ice by two rounds of sonication for 2 mins each with 10 second pulses and 15 second pauses at 50% amplitude using 435-C probe (Sonics Vibra Cell). Lysates were clarified by centrifugation at 20,000 x g for 45 mins at 4°C in a JA25.50 rotor (Beckman Coulter). Clarified lysates were applied to a gravity flow column several times over 2 ml Ni-NTA resin (Sigma) which was pre-equilibrated with 5 column volumes (CV) of lysis buffer. The resin was washed with 5 CV of lysis buffer containing 0-, 40- and 80-mM imidazole, pH8 (Sigma) to remove nonspecific binding of contaminants. Recombinant proteins were eluted off the resin in four fractions by passing 4 CV of the elution buffer (lysis buffer containing 400 mM imidazole, pH8 (Sigma). Eluates were resolved on NuPAGE Novex 4–12% Bis–Tris Protein gels (Life Technologies) and stained with Instant Blue (Novus Biotech) to assess the purity of the recombinant proteins. Eluted proteins were further purified over gel filtration at 4°C on Superdex 200 increase 10/300 GL (Cytiva) using lysis buffer A. Proteins were snap frozen in liquid nitrogen and stored at −80°C for further use.

### Purification of 3-headed ODA complexes from *Tetrahymena* cilia

Axonemal ODAs were purified using high salt extraction of cilia as previously described (Mali et al., 2021). Briefly, large scale (6L) *Tetrahymena* cultures grown in SPP medium containing glucose and FeCl3 were deciliated using dibucaine (0.3 mM). Isolated cilia were pelleted at 13,500 g at 4°C. Cilia pellets were washed in cilia isolation buffer [CIB: 20mMHEPES (pH 7.4), 100mM NaCl, 4mM MgCl2, 0.1 mM EDTA] and de-membranated with 0.25% Triton-X detergent, freshly added protease inhibitors, 1 mM DTT and 1 mM PMSF followed by a 30-min incubation on ice. Following a wash in CIB to remove excess Triton-X and another spin to pellet de-membranated axonemes, complexes bound to the axonemes were extracted by incubating the axoneme pellet for 30 min on ice in 3ml of a high salt buffer (20 mM HEPES (pH 7.4), 600mMNaCl, 4 mM MgCl_2_, 0.1mM EDTA, 1mM DTT, 0.1mM ATP, 1 mM PMSF). 0.5 ml of the high salt extract was loaded onto 6 identical 5-25% sucrose density gradients made in CIB and centrifuged for 16 hours at 33,000 rpm in an SW41 rotor (Beckman Coulter) at 4°C. Fractions containing ODA complexes (assessed by SDS-PAGE and gel staining) were pooled and further purified using a Capto HiRes Q 5/50 column. ODA complexes eluting at 300mM salt were inspected for purity and intactness (assessed by co-elution of heavy, intermediate and light chains) by SDS-PAGE and Instant Blue, Sypro ruby or Lumitein gel staining. ODA containing peak fractions were flash frozen in liquid nitrogen in 50% Glycerol or used freshly for reconstitution and displacement experiments.

### Purification of 2-headed ODAs from pig tracheal cilia and reconstitution with DNAAF9

Pig tracheal ODA complexes were purified following a previously described protocol^33^. Six pig tracheas were freshly acquired from a butcher and transported in ice-cold PBS. Tracheas were cleaned by removing excess connective tissue and fat. Cleaned trachea were further washed. Deciliation was performed using a buffer A (20mM Tris [pH 7.4], 50mM NaCl, 10mM CaCl_2_, 1mM EDTA, 7mM beta-mercaptoethanol, 0.1% Triton X-100, 1mM PMSF, MG132 and 4 Protease inhibitor tablets. Trachea were vigorously shaken in the de-ciliation buffer for a minute. The buffer was poured into 1L bottles and centrifuged multiple times to pellet cellular debris @ 1500xg for 2 minutes each time at 4°C. The clarified supernatant devoid of cellular debris was transferred into 250 ml bottles and centrifuged in in a JLA16.25 rotor (Beckman Coulter) at 12,000 x g for 5 minutes at 4°C to pellet cilia. The cilia pellet was resuspended in 25ml re-suspension buffer (20mM Tris [pH 7.4], 50mM KCl, 4mM MgSO4, 1mM DTT, 1mM EDTA, 1mM PMSF, 1 Protease inhibitor tablet) and centrifuged again to obtain a cleaner cilia pellet. A high salt extract containing ODAs and other axoneme bound proteins was obtained by resuspending the cilia pellet in 3ml Dynein Extraction Buffer (20mM Tris [pH 7.4], 600mM KCl, 4mM MgSO_4_, 1mM DTT, 1mM EDTA, 1mM PMSF, 0.1mM ATP) for 30 minutes on ice. The high salt extract was centrifuged at 31,000xg for 15 minutes and the supernatant was dialyzed into a lower salt dialysis buffer (20mM HEPES, 50mM NaCl, 4mM MgCl2, 1mM DTT, 0.1mM ATP, 1mM PMSF) for 2 hours. 50mM NaCl was used in the dialysis buffer to dialyse out KCl which impairs gradient fractionation. 0.5 ml of the dialysate was overlayed onto a 5-30% sucrose gradient made in dialysis buffer and resolved by centrifuging the gradients overnight (15-16h) at 168,000xg in a SW40 rotor (Beckman Coulter). The gradient was manually fractionated into 0.5 ml fractions and a small sample from each fraction was run on an SDS-PAGE gel and stained with Sypro Ruby to visualize protein bands. Fraction 14 contained high molecular weight bands which accumulated strong stain. To verify that these high molecular weight bands corresponded to ODA heavy chains, fraction 14 was subjected to mass spectrometric identification of all proteins co-fractionating in fraction 14 (**Fig S3, supplementary file 1**). Negative stain EM analysis was performed on fraction 14 to verify intactness of the purified ODA holocomplexes. Complexes in fraction 14 were also mixed with DNAAF9 and negatively stained to check the effect of DNAAF9 on ODA conformation compared to free ODA.

### Complex reconstitutions, size exclusion chromatography and Shulin displacement assay

Complex formation of Arl3^WT^, Arl3^Q70L^, Arl3^T30N^, or the ARL3-FYY triple mutant with Shulin or DNAAF9 was investigated by analytical size-exclusion chromatography using a Superdex 200 increase 10/300 column (Cytiva). 0.38 nmoles of Shulin or DNAAF9 proteins was incubated with 4-fold molar excess of Arl3 protein variants for 1 hr on ice. The mix was supplemented with 0.5 or 1 mM of GDP or GTP, applied to the size-exclusion chromatography column and eluted with one column volume of lysis buffer A. The elution profile was recorded and eluted fractions analysed by SDS-PAGE and stained with Instant Blue (Novus Biotech) to assess co-elution of the recombinant proteins as an indicator of complex formation between Shulin or DNAAF9 and the various Arl3 variants. For the Shulin displacement assay, purified ODAs were reconstituted with Shulin as described previously^8^. The reconstituted complex was then incubated with an excess of Arl3^Q70L^ in the presence of 1 mM GTP for 30 minutes on ice. The resulting mixture was resolved on a Superose 6 increase 10/300 column (Cytiva) and the eluted fractions were analysed by SDS-PAGE and SyproRuby or Lumitein staining.

### Negative-stain electron microscopy and data processing

Freshly isolated complexes between Shulin or DNAAF9 and Arl3 Q71L were diluted to between 0.01-0.05 mg/ml in gel filtration buffer and 3 μl of each sample was applied to freshly glow-discharged grids (300-mesh copper with formvar/carbon support, TAAB). Grids were glow-discharged using an Edwards Sputter Coater S150B or an EMS GloQube system for 15-30 s at 35 mA. Samples were incubated on grids at room temperature for 1 min and then blotted by wicking away excess liquid using a filter paper. 2% uranyl acetate was applied to grids for a minute and air-dried after wicking away excess liquid. Micrographs were acquired using FEI EPU on a FEI 200kV Tecnai 20 microscope equipped with either a FEI Ceta 4k x 4k charge-coupled device camera or a FEI Flacon II direct electron detector at a nominal magnification of 80,000x and a pixel size of 1.27 Å/pixel (DNAAF9-Arl3 complex) or 3.64 Å/pixel (Shulin-Arl3 complex).

Negative staining of free pig ODA holocomplexes and after combining with DNAAF9 was performed as described above. Micrographs were acquired at a nominal magnification of 30,000x and a pixel size of 3.803 Å/pixel on a JEOL 2100 Plus 200kV TEM equipped with a Gatan OneView camera. Small datasets were used to obtain representative 2D class averages of pig ODA alone (in an open conformation; 759 particles) and in presence of DNAAF9 (in a closed conformation; 740 particles).

All EM data processing was performed using RELION-3.2 or 4. For obtaining DNAAF9-Arl3 2D class averages and 3D map, first 1232 particles were manually picked from 7 micrographs. An inital 2D classification was performed to obtain class averages that were used to autopick 161,220 particles. Successive rounds of 2D classifications provided well-defined 2D class averages from 21,423 particles. Sub-classification of these particles yielded classes which contained DNAAF9 alone (13,444 particles) and DNAAF9 with an extra density corresponding to Arl3 (11,135 particles). The Shulin-Arl3 dataset was similarly processed to obtained 2D class averages of Shulin alone and Shulin bound by Arl3. The 11,135 particles from the DNAAF9-Arl3 dataset were used for 3D classification and refinement to obtain a low resolution (∼18 Å) map of the DNAAF9-Arl3 complex.

### AlphaFold (v2 and v3) monomer, multimer predictions and structural analyses

AlphaFold2 Multimer and monomer structural predictions were generated using the Google Colab Alphafold2 Notebook with a Colab Pro+ subscription^34, 35^. For multimer prediction, the algorithm was asked to model a 1:1 heteromeric complex between human DNAAF9 (UniProt ID: Q5TEA3) and ARL3 (UniProt ID: P36405). Default parameters were used for MSA generation, and the amber relaxation step was enabled. Monomer protein models for DNAAF9 and Shulin (UniProt ID: Q22YU3) in Fig. S1b and d were also generated using the Google Colab Alphafold2 Notebook. In each prediction, the highest ranked rank_1 model was downloaded in .pdb format for further analyses. Model statistics are as generated by the software with annotations to highlight relevant details. AlphaFold3^36^ (AlphaFold server: https://alphafoldserver.com/) was used for modeling larger complexes and presence of ligands. For predicting site 1 contact, a sequence comprising DNAH5’s helical bundles 5-7 was used with the full-length DNAAF9 sequence. For predicting site 2 contacts, a sequence containing DNAH5’s motor domain was used with the sequence for DNAAF9’s C1-C3 domains. Full-length DNAAF9 and ARL3 sequences were used to predict an interaction between the two in the presence of one molecule of GTP or GDP using the default parameters^36^. Domain level RMSD scores were obtained from the PDB using the tool for Pairwise Structural Alignment^37^ using the domain boundaries for Shulin and DNAAF9 as described in Mali et al. followed by confirmation of structural alignments in UCSF Chimera X^38^. The DNAAF9-ARL3 AlphaFold2 predicted model was rigid body fitted into the negative stain EM 3D reconstruction of the DNAAF9-Arl3 complex using UCSF ChimeraX.

### Bioinformatics and conservation analysis

Ortholog searches for human DNAAF9 and ARL3 were performed using PSI-BLAST. For DNAAF9, sequences from six representative model organisms with motile cilia (Pig, Mouse, *Xenopus*, Zebrafish, *Tetrahymena* and *Chlamydomonas*) were aligned using the tools embedded in UniProt (Clustal Omega)^39^. Sequence alignments were visualized using ESPript ^40^. Residue level conservation scores were calculated and plotted onto PDB structures using the CONSURF server for the predicted DNAAF9 monomer and the DNAAF9-ARL3 multimer complex^41^. Final visualisations were performed in ChimeraX

## Acknowledgements

The authors would like to thank Kazufumi Mochizuki for providing *Tetrahymena* strains, Ian Collinson, Ervin Fodor, Anthony Roberts and Georgia Isom for sharing critical equipment, the University of Bristol’s Proteomics and Wolfson Bioimaging Facilities for data acquisition. We thank Shona Murphy, Sumana Sanyal, Anthony Roberts, Katerina Toropova for critical reading of the manuscript and Clinton Lau for helpful comments. We also acknowledge the contributions of Rhiannon Hughes, Kincaid Ingram and Isaac Dowell in acquiring preliminary data, James Daly and Philip Lewis for advice on bioinformatic analyses of proteomics data. **Funding:** This study was supported by an MRC Career Development Award (MR/X007219/1), an Academy of Medical Sciences Springboard Award (SBF007\100151) and start-up support from the School of Biochemistry, Bristol and the Dunn School of Pathology, Oxford (EPA fund). **Author contributions:** K.H. b. I. purified all recombinant proteins and performed biochemical reconstitutions, displacement experiments and (with G.R.M.) performed bioinformatics, structural modeling and analyses. G.R.M. performed cellular studies and endogenous IPs for proteomics, conducted initial biochemical experiments, purified endogenous pig ODAs and (with K.H. b. I.) *Tetrahymena* ODAs. G.R.M. prepared negative-stain EM samples, acquired micrographs (with C.M.) and processed all EM data. With guidance from G.R.M., B.B. analysed cellular imaging data and M.R. performed endogenous IP-western blots. K.H. performed TMT labelling and acquired proteomics datasets which G.R.M. analysed and interpreted. G.R.M. conceived and guided the project. K.H. b. I. and G.R.M. prepared figures and wrote the manuscript with input from all the authors. **Competing interests:** The authors declare no competing interests. **Data availability:** The raw mass spectrometry proteomics dataset has been deposited to the ProteomeXchange Consortium via the PRIDE^42^ partner repository with the dataset identifier PXD052722.

